# Large-scale recovery of integron cassettes for gene discovery screens

**DOI:** 10.64898/2026.03.17.712381

**Authors:** Filipa Trigo da Roza, André Carvalho, Amalia Prieto, Paula Blanco, Ester Vergara, Rocío Lopéz-Igual, Modesto Redrejo-Rodríguez, Melanie Blokesch, José Antonio Escudero

## Abstract

Integrons capture and stockpile adaptive genes encoded in modular mobile genetic elements called integron cassettes (ICs). Present in 17% of bacterial genomes, integrons can harbor hundreds of cassettes, representing a hotspot of genetic variability. Some ICs encode antimicrobial or phage resistance genes, but most remain functionally uncharacterized, representing an untapped source of genes of biotechnological interest. Here, we present two tools, the cassette gatherer and hunter, that allow the swift establishment of libraries of genes either from genetically tractable strains or directly from DNA. By re-engineering a class 1 integron, these platforms capture single cassettes in a sequence and function-independent manner. When applied to *Vibrio* genomes, they recovered hundreds of single cassettes per assay with >99% specificity. To validate the usability of our tools for the discovery of new genes, we subjected these libraries to screens against phages ICP2 and T4, and identified nine phage-defense systems, including five previously undescribed. Hence, these tools provide a fast and simple method to recover thousands of ICs, in a way amenable to gene-discovery screens.

## INTRODUCTION

Integrons are bacterial genetic platforms that enable the acquisition of genes encoded in integron cassettes (ICs) through site-specific recombination^1,2^. ICs are stockpiled, forming a memory of functions that are adaptive for the host. Integrons were discovered for their role in disseminating antimicrobial resistance (AMR) genes among Gram-negative bacteria^3^ and have recently been shown to harbor phage defense systems^4–6^. However, integrons can capture and mobilize cassettes encoding genes with a broad variety of predicted functions beyond clinical relevance^7,8^.

These platforms can be divided into two main categories: mobile integrons (MI) and sedentary chromosomal integrons (SCI). MIs are plasmid-borne and associated with transposons, acting as vectors for more than 170 antimicrobial resistance genes and dozens of phage defense systems^4,5,9,10^. The class 1 is the most prevalent and best studied MI. It is abundant in the gut of humans and domesticated animals^11^, to the point where more than 10^23^ copies of integrons are released every day to the environment^12^. However, MIs represent only a small fraction of all integrons. SCIs are found in approximately 17% of bacterial genomes^13^ and constitute a much larger, underexplored, and underestimated reservoir of ICs. Most functions encoded in their cassettes are still unknown but are related to a wide variety of processes, including metabolism of lipids, carbohydrates, amino acids, and co-enzymes, among others^7,14,15^. Indeed, these elements can harbor hundreds of cassettes per cell, representing a hotspot of genetic variability in bacteria. For instance, the first SCI described was the superintegron (SI) of strain N16961 of *Vibrio cholerae,* where it spans 126 kb and carries 178 cassettes, accounting for ∼3% of the genome^16,17^; other strains, like W6G, carry 303 cassettes across 210 kb, representing 5% of the genome. The length and variability of SCI arrays underscore the role of integrons not just as vehicles for AMR, but as powerful, unexplored genetic reservoirs containing exaptable genes of adaptive value. Moreover, it is precisely from SCI arrays that MIs have captured cassettes that are valuable in clinical settings^18^. Understanding the functions encoded within SCIs is therefore also crucial for anticipating future adaptations mediated by integrons in pathogenic bacteria. From a structural perspective, integron platforms comprise three conserved elements: a gene encoding the integrase that governs all recombination reactions (*intI*)^19^, an *attI* site where incoming cassettes are inserted^20,21^, and a P_C_ promoter that drives their transcription^22–24^ (Fig. 1a). Integron cassettes typically contain a single open reading frame (ORF) and a recombination site known as *attC*^25^. A unique feature of *attC* sites is that they are highly variable. Their size ranges from less than 60 to more than 150 bp, and their sequence is not conserved except for the 5’-GTT-3’ triplet where the crossover takes place^26^. Instead, *attC*s show a conserved palindromic structure with important landmarks, like extrahelical bases, that allow for integrase recognition and adapt the recombination pathway^27–31^.

**Figure 1.**
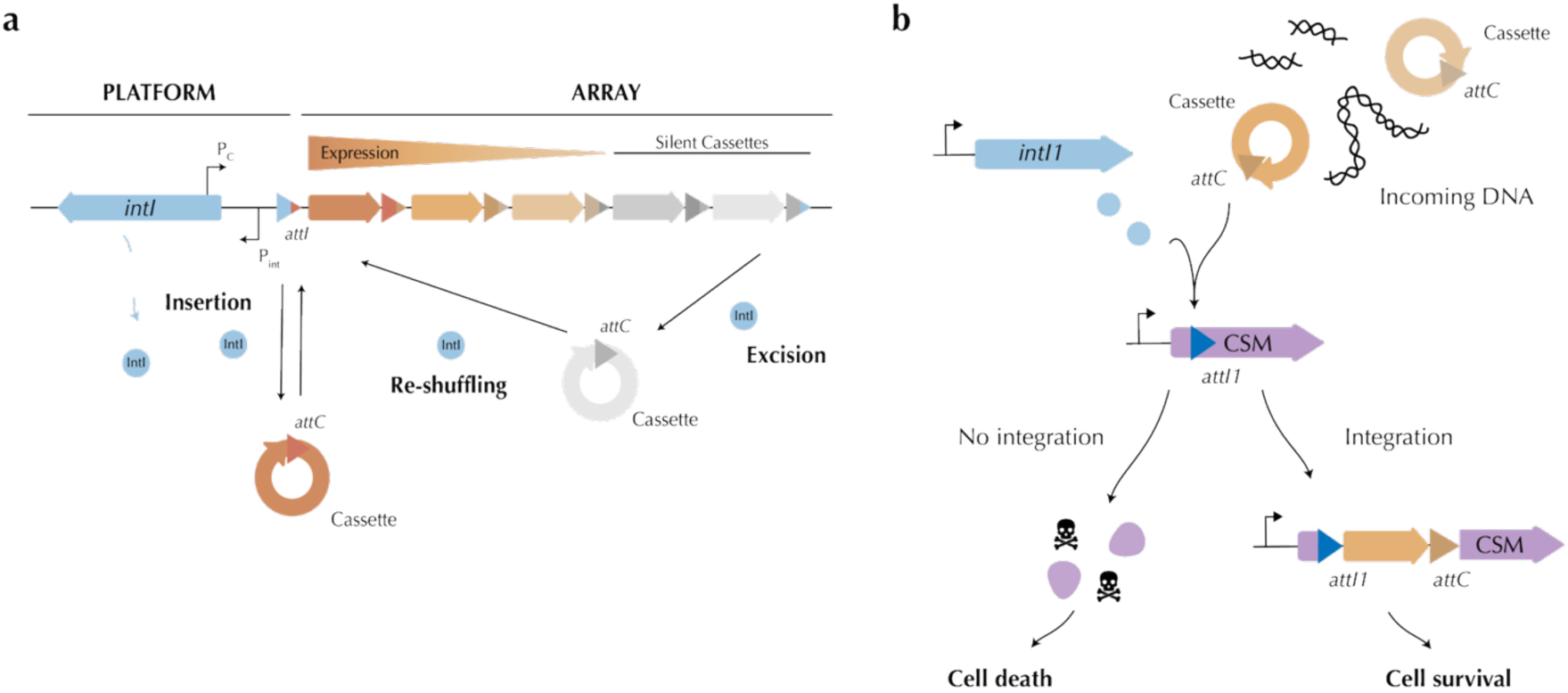
Leveraging the biotechnological potential of integrons for its "self"-detection. **a)** Schematic representation of an integron. The functional platform is depicted on the left, composed of the integrase-encoding gene (*intI*), the cassette promoter (P_C_), the integrase promoter (P_int_), and the integron insertion site (*attI*). The variable array of integron cassettes is represented on the right. Hybrid *attI* and *attC* sites are indicated with corresponding colors. The color intensity of each arrow reflects expression levels. The main reactions catalyzed by the IntI that lead to cassette insertion (*attI* × *attC*), excision (*attC* × *attC*), and reshuffling are also represented. **b)** Schematic representation of the cassette capture platform. We have re-engineered a class 1 integron to serve as a capture device using a counter-selectable marker (CSM) as a blind reporter. By embedding an integration site (*attI1*) within the CSM gene, incoming cassettes disrupt the CSM gene, and only recombinants survive.

Integron cassettes represent a remarkable biological resource, encoding extremely varied and adaptive functions^7,32,33^. For gene discovery, ICs offer a distinct advantage over conventional functional metagenomics: their intrinsic ability to be excised makes them independent of their genetic context, functioning as plug-and-play genes for their hosts^14^. Yet, access to functions encoded in cassettes and their exploitation has been hindered by the lack of sequence conservation of *attC*s. Indeed, some PCR-based methods have been developed, but they typically rely on primer binding to sequences that are either present in only a small subset of MIs^34–36^ (leaving most MIs and all SCIs unexplored), or within *attC* sites that are –by definition– not universally conserved^37,38^. Instead, producing synthetic gene libraries has recently emerged as a viable alternative, enabling the discovery of 60 phage defense systems^4,5^, as well as the in-depth characterization of AMR cassettes^10,39^. However, these approaches remain costly and time-consuming, limiting their application for routine purposes.

Here, we present two genetic tools that allow the high-throughput exploration and exploitation of integron cassettes. We have re-engineered a class 1 integron to generate libraries of single cassettes in a sequence, genetic context-, and function-independent manner. To do so, we have embedded the *attI1* insertion site into different counter-selectable marker genes (CSM), allowing cells to survive only if a cassette has been recruited (Fig. 1b). We have created a plasmid-based version of the tool – the cassette gatherer – that once transferred into the target host can rapidly (<24h) generate cassette libraries from SCIs with high efficiency (>99%) and specificity (>99.9%). To overcome host transformation barriers, we also developed the cassette hunter, which captures cassettes directly from DNA samples. For this, we established a modified version of the tool in the chromosome of the *hapR*^+^ strain of *Vibrio cholerae* N16961 ΔSI^14^, to take advantage of its natural transformation ability, while avoiding interference from SI cassettes. The cassette hunter is extremely versatile, recruiting hundreds of cassettes with high efficiency (89 to 97%) and specificity (>99.9%), providing only genomic DNA (gDNA) from the bacterium of interest. To demonstrate the utility of these tools for gene discovery, we built extensive libraries from multiple *Vibrio* spp. genomes and searched for novel phage defense systems. As a proof of concept, we challenged the libraries directly with Vibriophage ICP2, or indirectly, with phage T4 in *E. coli*, and successfully detected four known systems and five uncharacterized ones. Together, the cassette gatherer and hunter provide a sensitive and specific method to capture integron cassettes and exploit their biotechnological potential.

## RESULTS

### Development of a platform to select for integron recombination events

Classical experiments to detect integron cassette recombination events are common in laboratory conditions but are based on selecting for the phenotype encoded in the cassette^18,40^. Here, we sought to engineer a platform capable of reporting the integration of cassettes regardless of their phenotype. We reasoned that embedding an integration site (*attI*) within a counter-selectable marker (CSM), incoming cassettes should disrupt it, and only recombinant cells would survive (Fig. 1b). To capture cassettes, the integrase would be expressed from an inducible promoter in *trans*. Hence, an efficient platform should meet several criteria: the CSM::*attI* should kill efficiently only under certain conditions (that we refer to as killing conditions), but should otherwise be innocuous to the host; and the *attI* should be functional, allowing for the integrase-mediated capture of cassettes. Cassette insertion should, in all cases, disrupt the CSM.

To engineer this cassette capture platform, we chose to modify a class 1 integron because its integrase can recognize *attC* sites from very different phylogenetic origins^40^. We first developed a plasmid-based tool using the CcdB/CcdA toxin-antitoxin system from *Vibrio fischeri* as CSM. We embedded the *attI1* site in frame with the *ccdB* gene, within a flexible loop in CcdB^41^, where AlphaFold3^42^ predicted that the structure of CcdB was not affected (Fig. 2a and Extended Data Fig. 1a and 1b). To test the functionality of the CcdB*::attI1* toxin, we cloned it under the control of the P_Lac_ promoter on a plasmid also containing *intI1* controlled by the vanillate-inducible P_VanM_ promoter^43^, and we inserted it in an *E. coli* bearing a second plasmid encoding the *ccdA* antitoxin under the control of a P_BAD_ promoter (Fig. 2b). We calculated the killing efficiency of *ccdB::attI1* by plating cells in survival (inducing *ccdA*) and killing conditions (repressing *ccdA*). Survival rates (number of colony-forming units (CFU) in killing conditions over CFUs in survival conditions) were of 10^-6^, in line with previous reports using *ccdB* fusions^41^, and no significant differences were found with the WT *ccdB* allele compared to the *attI*-interrupted version (Fig. 2c). This shows that *ccdB::attI1* is functional and has a large dynamic range, spanning at least five orders of magnitude before retrieving false positives. These were originated by the disruption of *ccdB* mediated by insertion sequences (IS) (Extended Data Fig. 1c). Then, to test if cassette insertion would effectively impair CcdB-mediated toxicity, allowing for cell survival, we inserted a set of eight ICs of different sizes and functions into the *ccdB*-inserted *attI* site, mimicking cassette capture events. In all cases, *ccdB* was no longer active, leading to 100% cell survival (Extended Data Fig. 1d). Hence, our system can report cassette capture in a way agnostic to the function encoded.

**Figure 2.**
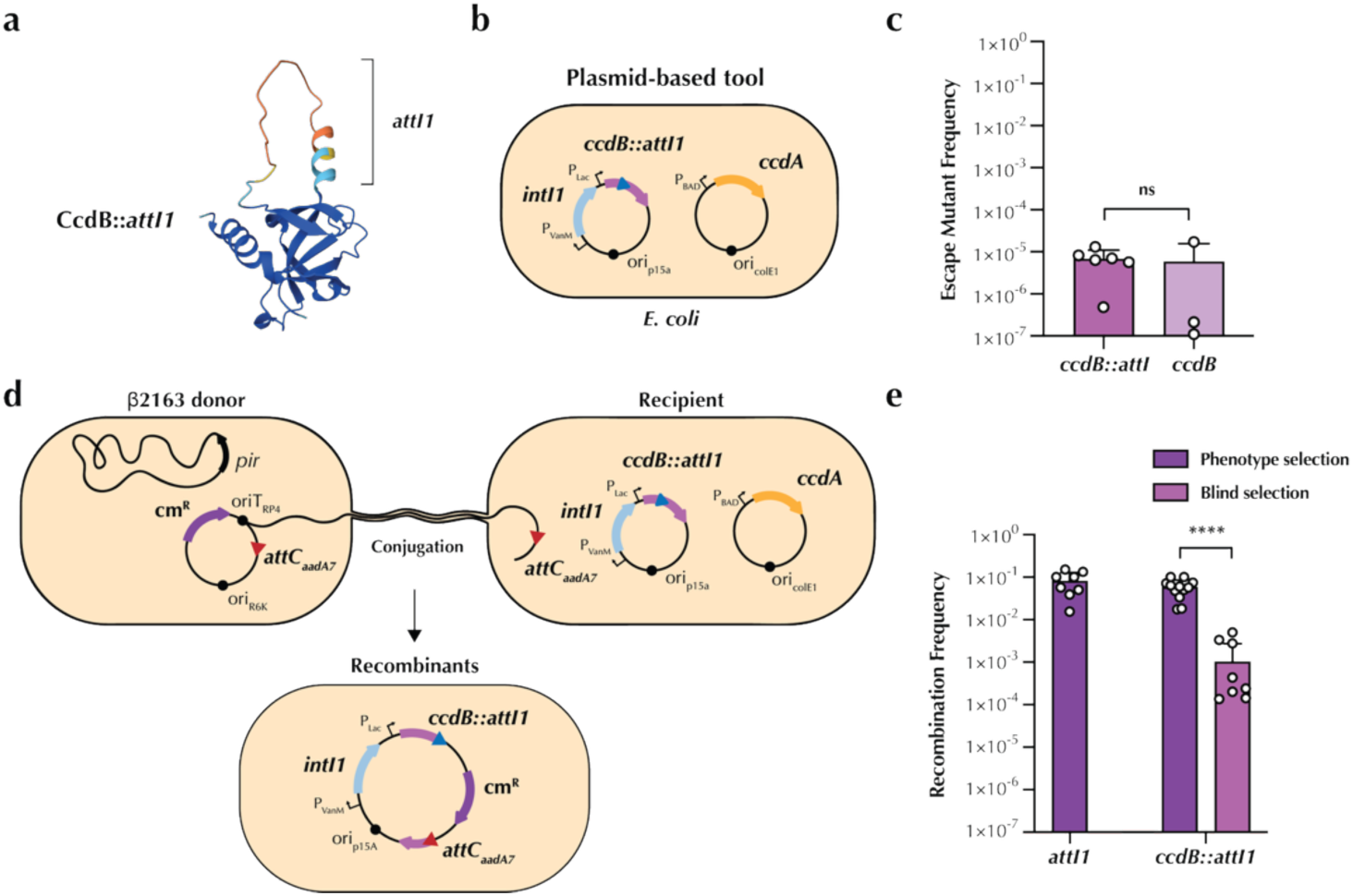
A plasmid-based tool to select for integron recombination events. **a)** Predicted structure of CcdB::*attI1* fusion protein, highlighting the translated *attI1* site. Models were generated using AlphaFold3^42^. Blue to red scale indicates high to low confidence in folding prediction. pTM = 0.78. **b)** Schematic representation of the IC capture platform in a plasmidic setup. **c)** Killing assay in *E. coli* comparing *ccdB::attI1* to the WT gene. Cells were grown in either survival conditions (with arabinose 0.2% that induces the expression of *ccdA*) or killing conditions (with glucose 1% that represses the expression of *ccdA*). The escape mutant frequency was calculated as the ratio of the CFUs growing in glucose and the total CFUs growing in arabinose. **d)** Schematic representation of the classical recombination assay. The suicide vector pSW23T carrying an *attC* site (red triangle) was delivered through conjugation from the *E. coli* β2163 donor strains to recipient strains containing the plasmid-based *ccdB::attI1* tool. Recombinants were selected either by phenotype, for chloramphenicol resistance on selective plates, or by blind selection, under *ccdB* expression. **e)** Recombination frequencies of the plasmid-based tool comparing phenotype selection and blind selection with the *attI1* control. Bar charts show the mean of the escape mutant frequency ± s.d. from biological replicates (individual dots). Statistical significance was determined using the Mann-Whitney U test; ns, not significant; ****, *p* < 0.0001.

We sought to confirm that the *attI1* was functional, allowing the platform to deliver the integrase-mediated capture and reporting, independently of the phenotype conferred by the cassette. To do this, we performed a well-known recombination assay in the field^40,44^ in which a suicide plasmid carrying an *attC* site and a chloramphenicol resistance (Cm^R^) gene is conjugated to our reporter cell. Then, the P_VanM_-induced integrase delivers the *attC* x *attI1* reaction (cassette capture) and the suicide plasmid is integrated into the integron array and therefore maintained (Fig. 2d). In our experimental setup, recombinants can be selected by the Cm^R^ phenotype—as it is normally done—or through blind selection, mediated by the killing activity of *ccdB*. Our results show that blind selection is possible at frequencies well above the background noise, suggesting that our aim is feasible. However, phenotypic selection was considerably more efficient (10^-1^ or 10^-2^) than blind selection (Fig. 2e). We interpret this as a phenomenon of genetic dominance consequence of using a multicopy plasmid^45^. In this scenario, non-recombined alleles of the CSM present in some plasmid copies would still kill the host, despite successful cassette capture in other copies. Despite this, we reasoned that this plasmid-based tool could be useful and potentially very efficient in a context where cassette abundance and replication are not limiting factors, increasing the chances of all copies of *ccdB::attI1* to be disrupted (see the cassette gatherer below).

Altogether, these experiments show that blind selection of cassette integration events is possible, setting the stage for a tool to detect ICs, regardless of the functions encoded by captured cassettes.

### The cassette gatherer – a tool to create IC libraries

Having successfully built a counter-selectable capture platform, we aimed to tackle a major challenge in the field of integrons: establishing cassette libraries from SCIs arrays. To do so, we targeted the SI of *V. cholerae* N16961 by introducing a modified version of the plasmid-based tool in this strain. Plasmid transformants were incubated for 4h inducing the expression of *intI1* and *ccdA*, and then plated in killing and survival conditions to recover recombinants and total cells, respectively (Fig. 3a). We obtained thousands of colonies in killing conditions, yielding recombination frequencies of 10^-3^, well above the escape mutant frequency of an *intI1*^-^ control (at 10^-6^) (Fig. 3b). We initially sequenced the inserts of 81 colonies to verify the capture events and found that all colonies contained a cassette from the SCI, totaling 65 different cassettes (Extended Data Fig. 2a and Extended Data Table 1). These results suggest that we can swiftly capture cassettes from SCIs, establishing large gene libraries.

**Figure 3.**
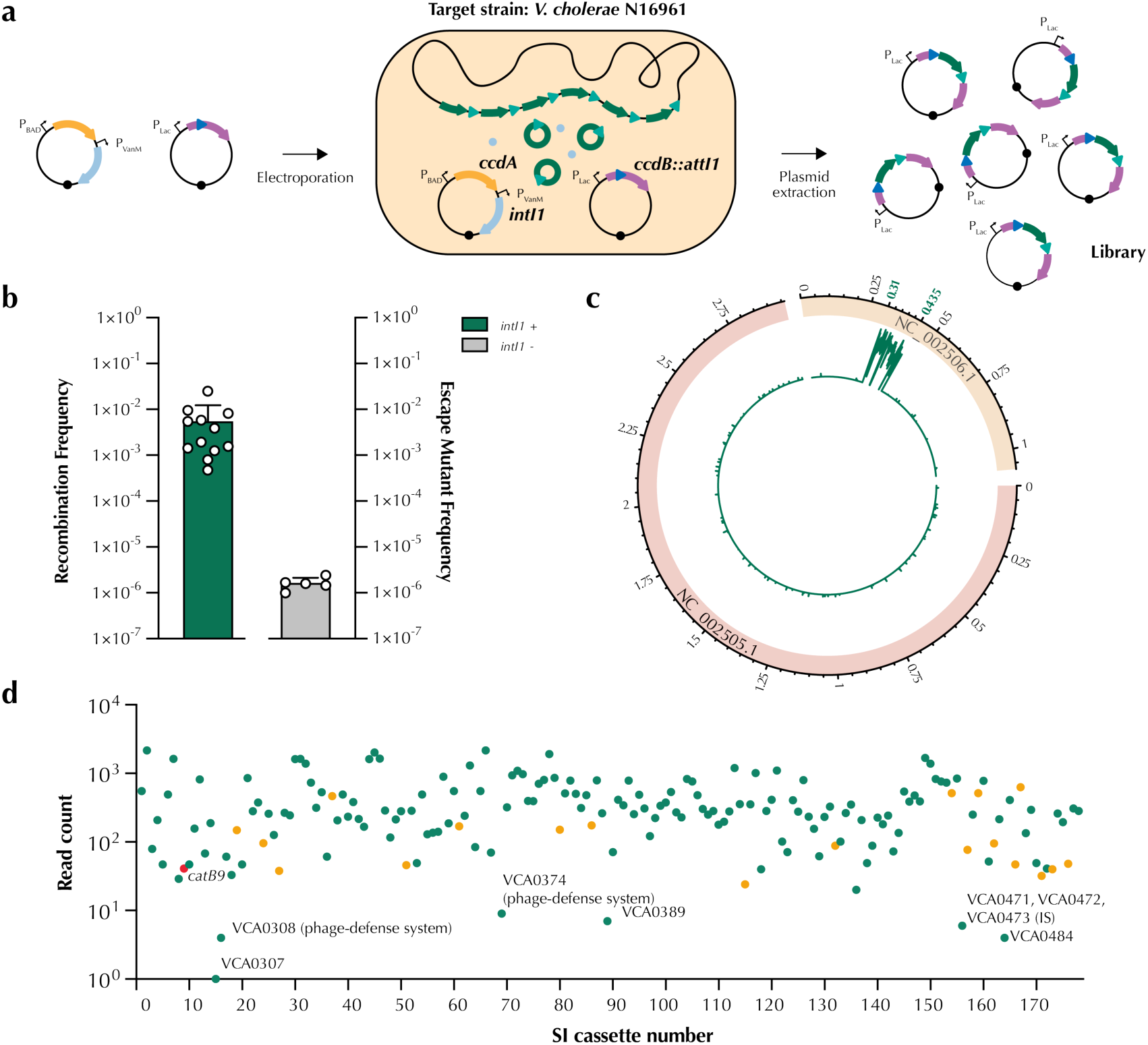
The cassette gatherer. **a)** Schematic representation of the library generation using the cassette gatherer. After the electroporation of the plasmids containing the *ccdB*-based tool into the targeted strain, we induce the expression of *intI1* in survival conditions (expressing *ccdA*). After growing cells for 4h in these conditions, cells are plated in killing conditions. Only cells that have captured cassettes in our *ccdB::attI1* bearing plasmid can survive. Plasmids are then extracted directly from the targeted strain. **b)** Recombination frequencies of the library generation from the *V. cholerae* N16961 strain. The escape mutant frequencies, depicted in grey (*intI1* -), correspond to a strain where we transformed a Δ*intI1* version of the *ccdA* plasmid. The recombination frequencies were calculated as the ratio of the number of recombinants and the total CFUs growing in arabinose. Bar charts show the mean of the recombination frequency ± s.d. from biological replicates (individual dots). **c)** Average coverage per kilobase (green plot) given in log10 reads of the two chromosomes of *V. cholerae* N16961 (RefSeq GCF_000006745.1). The sector NC_002505.1 corresponds to chromosome 1 and NC_002506.1 to chromosome 2. Chromosome coordinates are marked in the outside circle in black, and the superintegron coordinates are marked in green. **d)** Read count of the 178 cassette features. The *catB9* cassette is marked in red, and the 19 TA cassettes are marked in orange. Cassettes with lower (<10) read counts are labelled with their ORF ID and, if known, their identified function.

To better analyze the proportion of the SCI that was retrieved, we amplified inserts from ∼10^4^ colonies and sequenced them using long-read sequencing. We obtained ∼45,000 reads (250x coverage of the SI). Sequence analysis showed that >99.9% of reads mapped to the *V. cholerae* N16961 superintegron, demonstrating the high specificity of the method (Fig. 3c). The great majority of the reads mapped to only one cassette (95%) (Extended Data Fig. 2b), suggesting that our approach mainly recovers single cassettes. All cassettes from the SCI were represented in the library, with counts per cassette ranging from 1 to 2,000, with a mean of 419 (Fig. 3d and Extended Data Table 2). The only exception was cassette 68 (0 reads), which encodes the Sucellos phage defense system and contains an IS, locating the two *attC* sites 4 kbp apart. Cassette size is known to be important for recombination^46^, so we inferred that insertion of the IS makes Sucellos no longer mobile. Indeed, we found a negative correlation between cassette size and counts in our library (R=-0.54; *p*=1.3e-14), (Extended Data Fig. 2c). The least represented cassettes (1 to 10 reads) include other phage-defense systems (Ogmios and Divona), another long cassette containing an IS (VCA0471-3) and three cassettes of unknown function (Fig. 3d). Notably, TA systems were not among the least represented cassettes, highlighting that their movement is not toxic as long as the cassette is not lost.

Overall, we demonstrate that the cassette gatherer is a powerful tool capable of creating complete SCI libraries with high efficiency and specificity in less than 24 hours. It offers, for the first time, a reliable and high-throughput way to recover SCI-encoded genes.

### The cassette hunter – a tool to search for ICs directly from DNA samples

While the cassette gatherer significantly advances our ability to study cassettes from SCIs, it is limited by the need to introduce a plasmid in the target host. We hence deemed it essential to develop a tool capable of recovering cassettes directly from DNA, which would work on non-transformable microorganisms. *Vibrio cholerae*, renowned for advancing our understanding of chromosomal integrons^16^, is also a naturally competent bacterium^47^. This makes it an ideal candidate to establish a tool capable of detecting ICs from DNA samples. Since in this case the uptake of DNA can be limiting, we sought to free our tool of the negative effect of genetic dominance. For that, we developed a monocopy version of the tool, inserting it into the chromosome. Initial tests using the CcdB/CcdA system revealed a non-negligible level of toxicity. We hence chose the *sacB* gene from *Bacillus subtilis* as a CSM^48,49^. This gene encodes an enzyme that synthesizes the fructan polymer levan from sucrose, which can inhibit the growth of some Gram-negative bacteria, including *V. cholerae*. We inserted the *attI1* in three flexible linking regions that did not alter SacB folding (SacB::*attI1*_300_, SacB::*attI1*_203_, and SacB::*attI1*_403_) (Fig. 4a and Extended Data Fig. 3a and 3b). We inserted these fusions in the chromosome 1 of *V. cholerae* N16961 ΔSI *hapR*^+^ (see methods for more information on this strain), under the control of the constitutive promoter of *sacB* (Fig. 4b). To avoid the interference of our platform with cassettes from the superintegron, we used a ΔSI knockout strain^14^. The killing efficiency of SacB::*attI1*_300_ (hereafter referred to as SacB::*attI1*) under killing conditions (in presence of sucrose and absence of NaCl in LB medium) was highest (Extended Data Fig. 3c) and was therefore selected for further experiments. We then introduced the pBAD::*intI1* plasmid and performed a killing experiment. Our results show that SacB::*attI1* kills efficiently and that the escape mutant frequency is low (10^-6^) and comparable to the frequencies observed for the non-modified *sacB* version (Fig. 4c). In this version of the tool, the escape mutants corresponded mainly to INDELs in the *sacB* gene (Extended Data Fig. 3d). Importantly, killing conditions did not affect the growth of the WT strain lacking *sacB*, validating the killing range (Extended Data Fig. 3e). After this, we subjected this tool to the classical recombination assay to evaluate the functionality of the *attI1* site and the efficiency of blind selection (Fig. 4d). In this setting, recombination frequencies of blind selection were close to those of the phenotypic selection (10^-3^), suggesting we successfully avoided the dominance effect observed with the plasmid-based platform (Fig. 4e).

**Figure 4.**
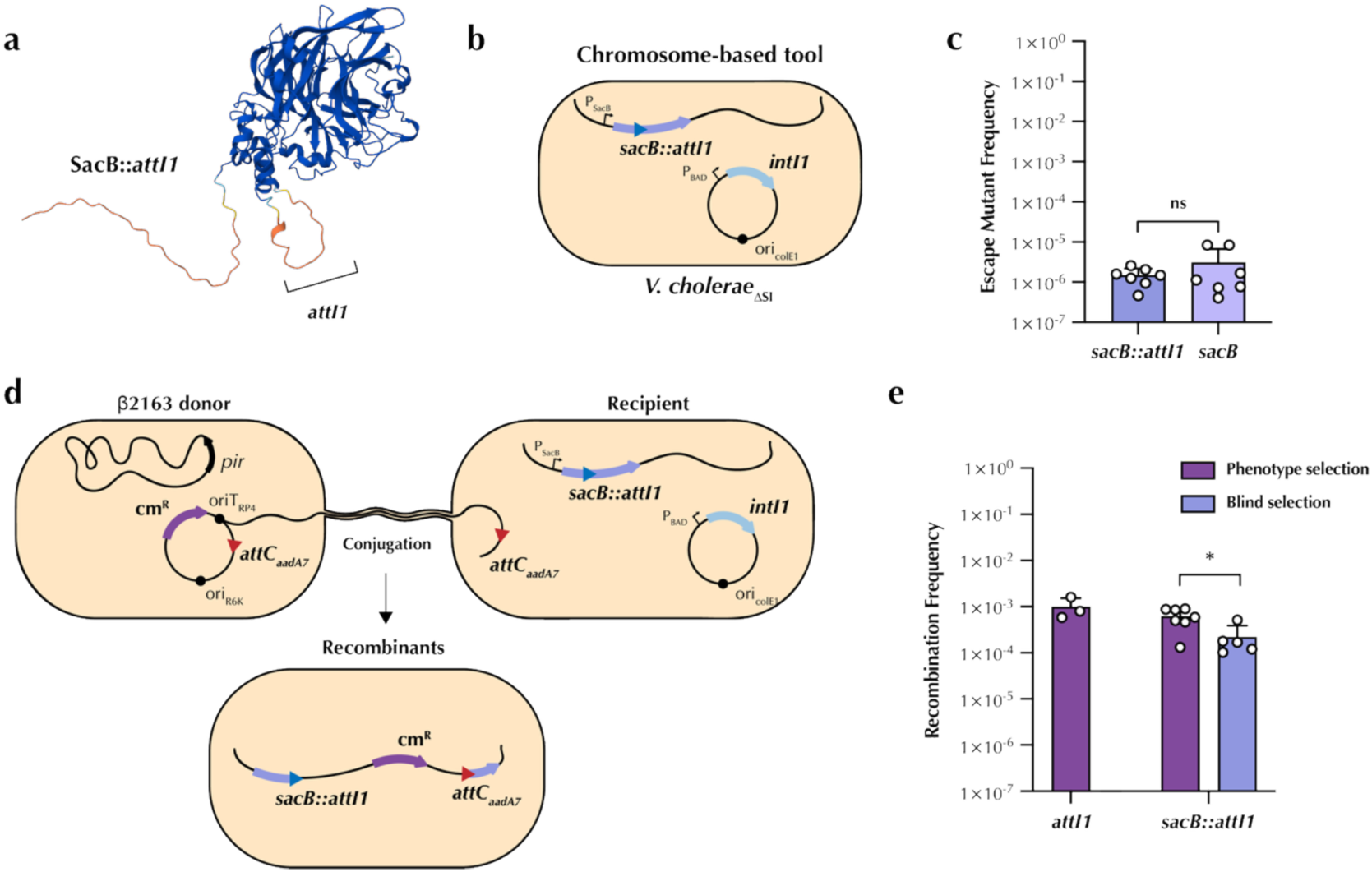
A chromosomal tool to select for integron recombination events. **a)** Predicted structure of the SacB::*attI1* fusion protein, highlighting the integration site and altered folding. Models were generated using AlphaFold3^42^. Blue to red scale indicates high to low confidence in folding prediction. pTM = 0.9. **b)** Schematic representation of the IC capture platform in a chromosomal setup. Transcription is granted by the P_SacB_ promoter. **c)** Killing assay performed in *V. cholerae* N16961 ΔSI comparing *sacB::attI1* to the WT gene. Cells were grown in either survival conditions (in LB medium) or killing conditions (in LB (∅NaCl) + sucrose 10%). The escape mutant frequency was calculated as the ratio of the CFUs growing in LB (∅NaCl) + sucrose 10% divided by the total number of CFUs grown in LB. **d)** Schematic representation of the classical recombination assay. The suicide vector pSW23T carrying an *attC* site (red triangle) was delivered through conjugation from the *E. coli* β2163 donor strains to recipient strains containing the chromosome-based *sacB::attI1* tool. Recombinants were selected either by phenotype, in chloramphenicol-containing plates, or by blind selection, in LB (∅NaCl) + sucrose 10%. **e)** Recombination frequencies of the chromosomal-based tool comparing phenotype and blind selection with the *attI1* control. Bar charts show the mean of the recombination frequency ± s.d. from biological replicates (individual dots). Statistical significance was determined using the Mann-Whitney U test; ns, not significant; *, *p* < 0.05.

We then attempted to improve the natural competence of the *V. cholerae* chassis and optimize it for integron recombination. Previous studies indicated that DNA uptake via natural transformation of a plasmid containing an *attC* site, followed by *intIA*-mediated cassette acquisition, yielded recombinant frequencies of approximately 10^-7^ ^50^. These frequencies can be too low to be distinguished from the background noise of spontaneous mutants in our setting (see Fig. 4c). We hence sought to increase this rate by generating and testing knock-out mutants in genes that could negatively affect the different parts of the process, such as the quorum sensor regulatory gene *luxO*^47,51^, the extracellular nuclease Dns^52^ that inhibits natural competence when not properly repressed, or the phage/plasmid defense systems *ddmABC* and *DE* that limit plasmid maintenance in pandemic *V. cholerae*^53^. We also tried providing the master regulator of natural competence in *trans*, *tfoX*^47^, controlled by the inducible P_BAD_ promoter. The combination of *tfoX* induction using a *Δddm* mutant allowed us to capture cassettes from DNA samples at significantly higher rates, with a 4-fold increase compared to the parental strain (Extended Data Fig. 4). We then introduced in this mutant the *sacB*-based platform, creating our final tool: the cassette hunter. After decoupled induction *tfoX* and *intI1* (see methods), cells were washed and incubated overnight in defined artificial seawater (DASW) with chitin to induce full competence, adding 4 µg of the tested gDNA. Finally, cells were plated in survival and killing conditions to recover recombinants (Fig. 5a). The system was validated by testing a PCR-amplified sample containing a two-cassette array, yielding thousands of colonies and recombination rates of 10^-3^. This exceeded escape mutant frequency by >100-fold and was comparable to homologous recombination rates, suggesting that integron capture was very efficient (Fig. 5b). We then tested gDNA samples obtained from six *Vibrio* spp. strains encoding SCIs and belonging to five different species (*V. cholerae* N16961, *V. cholerae* SA5Y, *V. cholerae* W6G, *V. vulnificus* NBRC 15645, *V. mimicus* MB461, *V. parahaemolyticus* RIMD 2210633, and *V. fortis* LMG 21557). Again, recombination rates exceeded the detection threshold by at least 100-fold in most cases and the efficiency of recombination was high enough to yield thousands of colonies in a standard assay (Fig. 5c). Notably, the recombination frequency over the total number of cassettes in the SCI of the tested gDNA was similar in all cases (Extended Data Fig. 5a); for instance, DNA of *V. fortis*, which carries only five cassettes, yielded the lowest recombination frequencies (10^-5^) while that of *V. cholerae* W6G, with 303 cassettes, showed frequencies of almost 10^-3^. We then sequenced these libraries using long-read sequencing after PCR amplification. We found that the cassette hunter recovered 89 to 97% of the cassettes present in the SCI of the sample, enabling the generation of libraries from large chromosomal integrons (Extended Data Tables 3-8). This tool also showed an excellent specificity for integron cassettes, as >99.9% of the reads mapped to the SCIs (Extended Data Fig. 5b). We compared the results generated with the cassette hunter and the cassette gatherer when taking cassettes from the genome of *V. cholerae* N16961 and found that cassette counts correlate significantly, validating the reproducibility of our results with different tools (Extended Data Fig. 5c). As before, the majority of the reads (<92%) corresponded to a single IC capture event (Extended Data Fig. 5d). Additionally, the library obtained with the cassette hunter showed a significant correlation between cassette size and recovery rate, similarly to what we observed with the cassette gatherer (Extended Data Fig. 5e).

**Figure 5.**
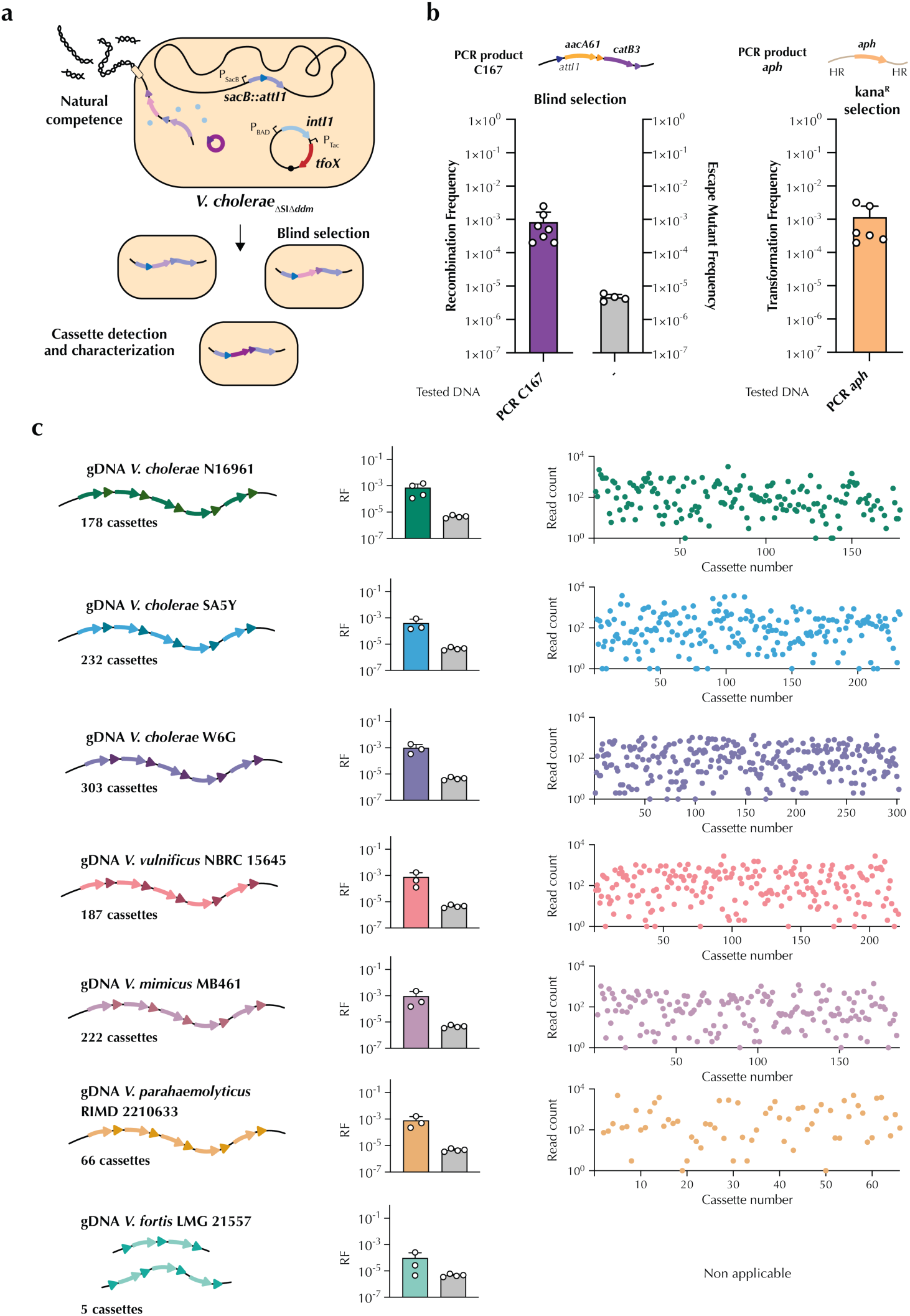
The cassette hunter. **a)** Schematic representation of the setup used to create the cassette hunter. In this system, the tested DNA sample is taken up by natural competence —upon induction of *tfoX*, the master regulator of competence—inside the *V. cholerae* ΔSI Δ*ddm* mutant, which contains a SacB-based version of our tool. The P_BAD_-induced IntI1 can mobilize the integron cassettes inside the *sacB::attI1* platform. Selection of recombinants is achieved through growth in killing conditions. **b)** Initial validation of the cassette hunter, testing a PCR product of an integron containing two cassettes compared to two separate controls: a reaction with no DNA (measuring the escape mutant frequency) and a sample testing natural competence and homologous recombination (PCR *aph* sample) by plating in kanamycin. Recombination and transformation frequencies were calculated as the ratio of the number of recombinants/transformants and the total CFUs. **c)** Recombination frequencies (RF) and library analysis of the tested gDNA from the sedentary chromosomal integron (SCI)-containing Vibrios. RFs were calculated as stated above and are compared with the negative control (escape mutant frequency). Library analysis is represented by the read count retrieved for each cassette feature of the SCI of the corresponding strain. Note that for *V. fortis,* the analysis was not applicable, as it contains only 5 integron cassettes in its SCI.

With these experiments, we show that the cassette hunter is capable of detecting integron cassettes directly from DNA samples with high specificity and sensitivity, allowing the generation of large libraries of cassettes from SCIs.

### Unveiling the genetic potential of integrons

To demonstrate the applicability of our tools for discovering novel cassette-encoded functions, we explored whether our SCI libraries could reveal new and biologically relevant adaptive traits. As a proof of concept, we focused on phage-defense systems, which have recently been identified as components of integron platforms.

We screened our libraries with two complementary strategies (Fig. 6a). We first tested them directly, subjecting the library obtained with the cassette hunter chassis (*V. cholerae* N16961 *hapR*+ ΔSI) against the ICP2 vibriophage, to investigate whether phage-defense activity could be identified without intermediate cloning steps. As a second approach, we implemented an indirect strategy in which the cassette contents of the six libraries from *Vibrio* spp. genomes were amplified and cloned in pMBA, a vector mimicking the genetic environment of a class 1 integron where cassettes are inserted at the *attI1*, and are under the control of a strong Pc promoter (PcS)^10,22,39^. We introduced this library in *E. coli* MG1655 and challenged it with phage T4. This approach aimed to prove the mobility of the library to other species using a system granting high expression levels, in contrast with the weak expression from P_SacB_ in the original platform (Extended Data Fig. 6a).

**Figure 6.**
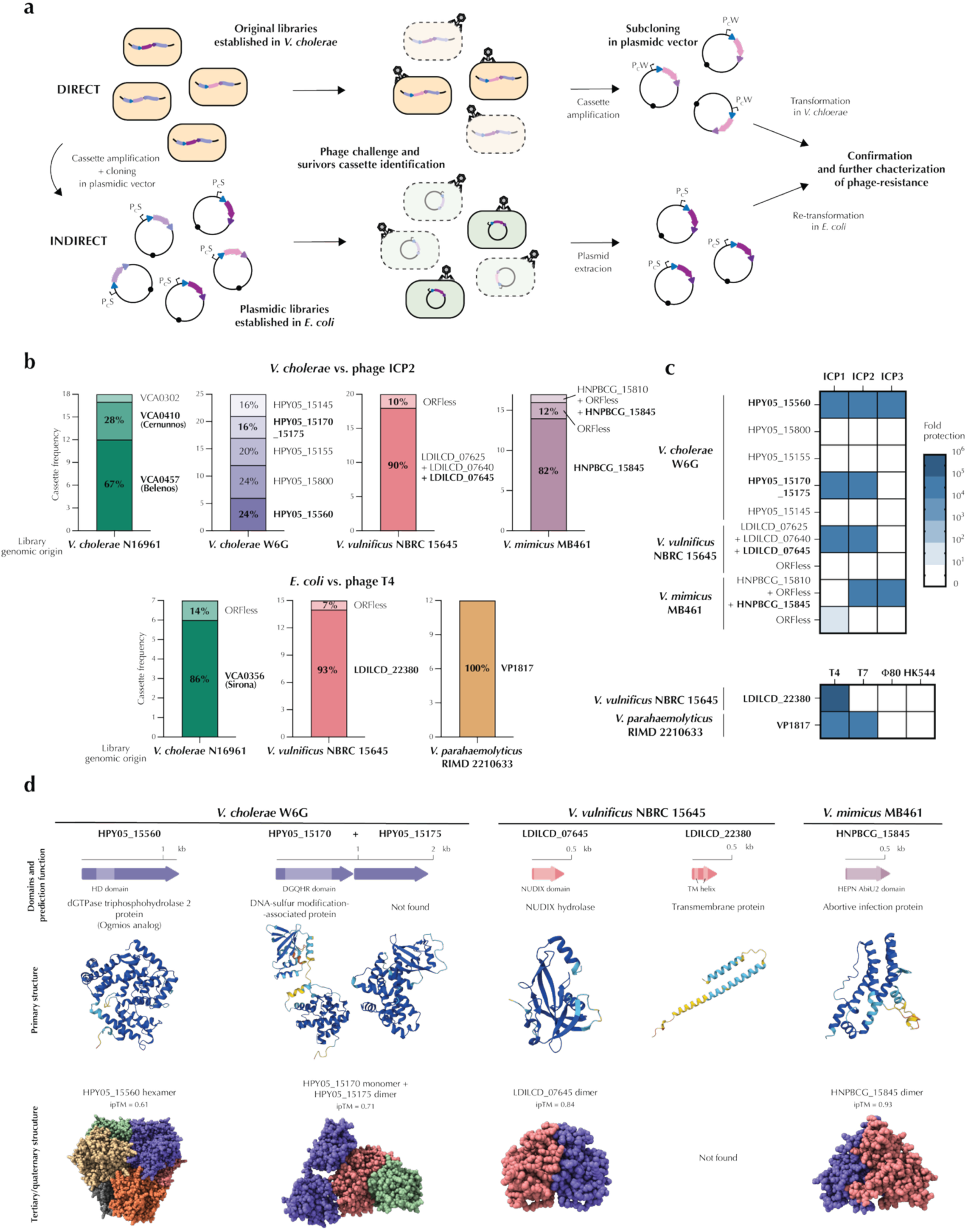
Screening for phage-defense systems. **a)** Schematic representation of the library screening workflow. Libraries were screened using two complementary approaches: (i) directly in the cassette hunter chassis using vibriophage ICP2, as previously described^54^. In this case, cassettes conferring survival were identified by Sanger sequencing and subcloned into a weak expression vector (P_C_W promoter) for phenotype verification; or (ii) cassette content was amplified and cloned into a strong expression vector (P_C_S promoter), introduced into *E. coli* MG1655, and challenged with phage T4 using the same strategy. Surviving clones were recovered, sequenced, and plasmids re-transformed for phenotype validation. **b)** Integron cassettes identified by Sanger sequencing and their corresponding frequency for each library showing survivors after phage challenge. **c)** Validation and characterization of phage resistance phenotypes. Candidate genes were subcloned in the pMBA vector and subjected to plaque assays. Heatmaps represent the mean fold resistance of three independent biological replicates against each tested phage. **b)** Genetic organization, predicted protein domains and function, and structural models of newly identified phage defense proteins. Protein structures were predicted using AlphaFold3^42^ and visualized with ChimeraX. Blue to red scale indicates high to low confidence in folding prediction.

To challenge these libraries, we employed a method known as “tab” (T4 abortive)^54,55^, which allows the recovery of phage-defense systems conferring resistance through direct protection, but also—and crucially—through an abortive infection mechanism (for more details, see Methods and^54^). We recovered colonies surviving ICP2 infection from the SCI libraries obtained with the genomes of *V. cholerae* N16961, *V. cholerae* W6G, *V. vulnificus* NBRC 15645, and *V. mimicus* MB461. Using phage T4, we recovered survivors from *V. cholerae* N16961, *V. vulnificus* NBRC 15645, and *V. parahaemolyticus* RIMD 2210633 SCI libraries (Fig. 6a). In all cases, we amplified and sequenced the cassettes encoded in a subset of colonies. Our results revealed four recently characterized phage-defense systems encoded in ICs: Cernunnos, Belenos, and Sirona from *V. cholerae*^5^, and VP1817 from *V. parahaemolyticus*^6^. Strikingly, we also recovered 14 ICs of unknown function (Fig. 6b). We subcloned cassettes found more than once in a clean genetic background and confirmed their resistance phenotypes (see Fig. 6a and Extended Data Fig. 6a). We failed to clone the IC containing HNPBCG_15845 alone, but successfully cloned the combination of cassettes that contains it (HNPBCG_15810 + ORFless + HNPBCG_15845). We then performed plaque assays against the vibriophages ICP1, ICP2, and ICP3, or the *E. coli* phages T4, T7, Φ80, and HK544 (Fig. 6c and Extended Data Fig. 6a and b). Our results showed that five cassettes conferred an increase in fold protection to the phage used for the challenge, and some conferred protection to three different phages. Interestingly, we also found that the recently described VP1817 system confers protection to T4 and T7 phages. This system was known to provide defense to phages Bas37, 38, and 40 from the BASEL collection^56^, but not to T4 and T7. Importantly, cassettes that did not confer protection were not prevalent in the screen (around ∼15% of false positives). The only exception was in the case of W6G, where they added up to 60%, suggesting that in some cases a second challenge could be useful to purify the library (see Fig 6b). Finally, when two or more cassettes had been captured by a single cell, we assumed that the phage-defense system was either the most prevalent in other combinations (as in the case of HNPBCG_15845, which was found in 94% of the surviving colonies from the *V. mimicus* library) or using additional analysis tools to predict protein functions and domains (as for LDILCD_07645 below).

We further characterized the phage-defense systems discovered here by searching for known functions and domains and predicting the protein structure (Fig. 6c). In *V. cholerae* W6G, DefenseFinder^57^ and PADLOC^58^predicted that cassette HPY05_15560 was a homologue of Ogmios^5^. This was the only cassette in our search where defense systems were predicted by algorithms. It encodes a protein containing an HD domain that seems to form homo-hexamers and confers resistance to ICP1, 2, and 3. In the same library, we identified a cassette containing two ORFs, HPY05_15170 and HPY05_15175, that confer resistance to ICP1 and 2. The first ORF is a monomer encoding a DNA-sulfur modification-associated protein, and the second is a dimer with no homologs identified. In *V. vulnificus* NBRC 15655, gene LDILCD_07645 confers resistance to ICP1, 2, and 3 and was identified as a dimeric NUDIX hydrolase. In the same library, LDILCD_22380 encodes a small transmembrane protein conferring resistance to T4. Finally, in *V. mimicus* MB461, HNPBCG_15845 forms a dimer with an HEPN AbiU2 domain, suggesting that it confers resistance to T4 and T7 through an abortive infection mechanism.

Our results show that the genetic libraries generated with the cassette hunter are amenable to gene discovery screens either directly or in the heterologous genetic background of the experimenter’s choice. Well-designed screens allow detection of functions even in situations where direct positive selection is infeasible.

## DISCUSSION

In this work, we introduce a tool capable of detecting integron cassettes and generating libraries from large chromosomal integrons, unlocking the possibility of exploring the hidden functional potential of thousands of genes encoded in these genetic platforms^32^.

Compared with current methods, our tools combine high sensitivity and specificity with simplicity of use. This has major advantages for detecting integron cassettes, compared to previous approaches. PCR-based methods have been widely used, relying on primer binding to conserved sequences. For class 1 MIs, targeting what was called the 3’ conserved sequence (the *qacEΔsul1* cassette), was used with clinical isolates for many years. Additionally, when studying cassettes in environmental samples, degenerate primers have been designed to bind a variety of *attC* sites. Both approaches have strong biases^34–36^: the advent of genome sequencing has revealed that *qacEΔsul1* is present only in a limited number of class 1 integrons and absent in all other MI classes. As for the amplification of *attC* sites, these are intrinsically variable, and even using degenerate primers introduces a clear bias towards a specific group of sites. Additionally, because these primers bind to the *attC* site in a way that the crossover points are not included in the sequence, it is impossible to confirm that the amplification product is a *bona fide* integron cassette by testing the *attC* recombination site functionality. Other methods have been developed that harness the integron machinery to avoid the specific binding of primers. Rowe-Magnus and colleagues developed a three-plasmid method, based on the generation of fosmid libraries from DNA samples that were then recombined by an integrase onto a conjugative plasmid. This plasmid was then transferred into a recipient strain where its content could be studied^59^. Although effective, this strategy is labor-intensive and retrieves large cassette arrays rather than single units. In contrast, our system is fast; it does not require previous steps other than DNA purification and recovers a larger number of cassettes in a very specific manner. Additionally, in most cases, cassettes are captured as single units.

Our tools exploit the high specificity of integron recombination provided by the peculiarities of *attC* sites that are recombined as folded ssDNA molecules. By embedding an *attI* site within a counter-selectable marker (CSM), cassette insertion disrupts the CSM gene, enabling blind selection of authentic ICs and preserving their natural *attC* boundaries (Fig. 2 and 4). This adaptation of a well-known molecular biology principle—interrupting a toxic gene to select for an insertion, as in the *ccdB*-based pTOPO system—had not previously been applied to integron recombination. Here, we show that *attI* insertion into CSM genes does not impair the toxicity of the fusion protein and that the approach efficiently recovers single, intact cassettes. This, together with other pioneering studies in the field of integrons^60,61^, highlights the biotechnological potential of these platforms *per se*.

Both the plasmid-based cassette gatherer and the chromosomal cassette hunter yielded highly efficient library generation, with >10^4^ and >10³ independent clones, respectively, and with negligible background (∼0.1-1%) (Fig. 3 and 5). Moreover, with the cassette hunter, escape mutants can be eliminated during sub-cloning in plasmidic libraries, since the amplification of non-recombined clones yields fragments below 30 bp that are lost during column purification of PCR products. Perhaps not surprisingly, the analysis of library composition revealed variation in cassette abundance, with some cassettes >1,000-fold more frequent than others. This likely reflects multiple factors: (i) size-dependent excision efficiency—smaller cassettes may impose fewer structural constraints between *attC* sites and thus be excised more frequently^46^; (ii) PCR amplification bias toward shorter fragments before sequencing; and (iii) toxicity of certain genes, which may reduce recovery of their cassettes. Supporting the first hypothesis, 95% of reads corresponded to single cassettes, suggesting that excision of two adjacent units is rare and that integron recombination naturally favors single-cassette capture. Indeed, in the cases in which we retrieved more than one cassette, these were non-contiguous in the DNA used. These tools have different advantages: the cassette gatherer is extremely efficient, retrieving all cassettes from an SCI (within the biological limits of the integron, i.e., cassette size). The cassette hunter allows for the testing of more variable microorganisms by using DNA directly as an input, albeit with a small loss of efficiency.

The scale recovery of thousands of genes and direct testing of generated libraries in just a few experiments offers substantial biotechnological potential. These collections provide a large source of uncharacterized genes for functional genomics, synthetic biology, and microbial engineering. As a proof of concept, we have used them to explore the presence of phage-defense systems. We have challenged our libraries with phages for which defense systems had been characterized, serving as controls. We retrieved most known systems while discovering five unknown ones, four of them with no homologs in DefenseFinder^57^ nor PADLOC^58^ (Fig. 6). This is consistent with recent reports highlighting integrons as reservoirs of such functions^4–6^. Moreover, thanks to the methodology developed by Vassallo *et al.*^54^, we were able to recover abortive defense systems that cannot be selected positively. This proves that any sound screen will be able to select cassettes among the thousands of options in our libraries. By decoupling cassette recovery from phenotype, our tools —the cassette gatherer and hunter— open the door to a high-resolution exploration of the vast and largely untapped genetic reservoir embedded in integrons.

## MATERIAL AND METHODS

### Strains, plasmids and primers

All strains, plasmids, and primers used in this study are listed in Supplementary Table 1.

### Growth conditions

*E. coli* strains were grown in LB Lennox plates (BD, France) supplemented with agar at 1.5% (Condalab, Spain) and incubated at 37°C unless stated otherwise. *V. cholerae* strains were grown in LB Miller agar plates (Sigma-Aldrich, USA) supplemented with agar at 1.5% (Condalab, Spain) at 30°C unless stated otherwise. Liquid bacterial cultures were incubated at 37°C/30°C with continuous shaking at 200 rpm. If necessary, the medium was supplemented with the corresponding antibiotic for the selection of the bacterial strain of interest. All the antibiotic and reagent references and concentrations used are listed in Supplementary Table 1.

### Plasmid constructions

Plasmids were assembled using an in-house-made Gibson Assembly^62^ or Fast-Digest restriction enzymes (ThermoFisher Scientific). For Gibson Assembly, primers were designed with 20-bp complementary ends for hybridization, and DNA fragments were synthesized if needed (IDT, USA). PCRs were performed using Phusion Green High-Fidelity Master Mix (ThermoFisher), and purified with GeneJet or High-Pure PCR kits (Roche). Digested fragments were ligated using either Gibson Assembly enzymes (50°C, 30 min) or T4 ligase (22°C, 2h). Cloning mixes were transferred into *E. coli* DH5α, *E. coli* MG1655, or *E. coli* π3813 (for *ccdB* constructs). Constructs were verified by colony PCR (DreamTaq, ThermoFisher) and Sanger sequencing (Macrogen, South Korea; Eurofins, France). Plasmids were purified (GeneJet Miniprep Kit) and electroporated into recipient strains.

### Chromosome Constructions

#### V. cholerae

Chromosomal insertions in *V. cholerae* were performed by natural transformation as previously described^63^ with modifications. Briefly, overnight cultures were diluted 1:100, grown to OD600 = 1.0, and incubated with chitin (150 µL slurry in DASW medium) at 30°C, in static conditions for ∼18h. Supernatant (550 µL) was removed, and purified tDNA was added for another ∼18h incubation. When necessary, reactions were outgrown in LB before selection on antibiotic plates. Constructs were verified by colony PCR and Sanger sequencing.

### Clean mutant generation in *V. cholerae*

Gene deletions were performed using suicide vectors (pGP704-Sac28)^64^, integrating via biparental mating followed by double recombination and sucrose-based counter-selection in NaCl-free LB medium. Transconjugants were plated on TCBS supplemented with ampicillin to confirm integration. Subsequently, colonies were grown in the absence of antibiotics to allow excision and loss of the suicide plasmid, which was verified by sucrose counterselection (LB ∅NaCl, 10% sucrose, 20°C). Deletions were confirmed by colony PCR.

### Killing assays

Killing assays were performed to (i) assess the efficiency of the CSM::*attI1* fusion proteins, (ii) establish the limit of detection of the generated tool, and iii) characterize the escape mutants of the system. Briefly, cells were grown in a liquid medium overnight in survival conditions, i.e., in the case of the CcdB/CcdA TA system, in the presence of the appropriate reagents to induce the antitoxin gene *ccdA* and repress *ccdB*. In the case of the *sacB* counter-selectable marker, bacteria were grown in standard LB medium. The next day, cells were diluted up to 10^-7^ and plated in 5 µL spots both in survival conditions, as previously described, or killing conditions. In the latter, for the CcdB/CcdA TA system, cells were plated in the presence of the appropriate reagents to repress the expression of *ccdA* and induce *ccdB*. In the case of the *sacB* counter-selectable marker, cells were plated in LB (∅NaCl) + sucrose 10% medium at 20°C. If fewer than 10 colonies were detected in the direct dilution spot, 100 µL of the overnight cultures was also plated in killing conditions. After this, the number of survivors in killing conditions over the total colony-forming units (CFU) was calculated, i.e., **escape mutant frequency**. Colony-PCRs were performed to characterize the escape mutants, and PCR products were sequenced by Sanger sequencing (Macrogen, Korea).

### Classical recombination assays

Classical recombination assays were performed to measure the efficiency of the *csm::.attI1* constructions in integron recombination. These experiments were essentially performed as previously described^40,44^ with some modifications. This assay simulates the natural conditions of cassette delivery through horizontal gene transfer by conjugating a suicide vector (pA123) that contains the *attC_aadA7_* site. This plasmid carries an RP4 origin of transfer (*oriT_RP4_*) oriented in such a way as to deliver the bottom strand of the *attC* recombination site. This ensures the transfer of the *attC* site in its recombinogenic form, thus facilitating integron recombination. The recipient strain cannot sustain the replication of the suicide plasmid; therefore, the only way for the vector pA123 to persist in the recipient cells is to recombine with the *attI1* site contained in the recipient strain. The cassette capture efficiency is measured by calculating the recombination frequency.

Briefly, the donor strain A123 was cultured overnight in LB medium supplemented with chloramphenicol 25 µg/mL and diaminopimelic acid (DAP) 0.3 mM. The recipient strain was cultured overnight in LB medium supplemented with appropriate antibiotics for correct selection and reagents for integrase repression. The next day, the donor culture was diluted 1:100 in LB with chloramphenicol and DAP, as indicated above, and the recipient strains were diluted 1:100 in LB with the same antibiotics and the appropriate reagents to induce *intI1*. These cultures were incubated until an OD_600_ of 0.7. Next, after washing the cultures, the recipient and donor were mixed in a total volume of 1 mL at a ratio of 4:1, centrifuged for 2 min at 8,000 rpm, and resuspended in 100 μL of LB. This volume was spread on a conjugation membrane (Millipore mixed cellulose ester membrane, 47 mm diameter, 0.45 μm pore size) on LB agar + DAP 0.3 mM + the appropriate reagent to induce *intI1*. Plates were incubated overnight (∼18h) to allow conjugation and recombination to occur. The following day, cells were resuspended in 5 mL of LB, after which 1:10 serial dilutions up to 10^-7^ were performed. 5 µL of each dilution was plated on LB medium supplemented with the appropriate antibiotics and reagents to recover the total recipient bacteria, and, to select for recombinant bacterial cells, dilutions were plated on LB medium with chloramphenicol either 25 or 2.5 µg/mL (for *E. coli* or *V. cholerae,* respectively), as well as other antibiotics and reagents needed. The ratio between recombinants growing in chloramphenicol and the total recipients was calculated, i.e., the <u>recombination frequency by phenotype selection</u>.

To count recombinant bacteria by survival from insertion within the chosen counter-selectable marker, cells were either plated in the appropriate reagents to induce the expression of *ccdB* and repress the expression of *ccdA*, or, in the case of *sacB*, cells were plated in LB medium (∅NaCl) + sucrose 10%, and incubated at 20°C. The ratio between the number of bacteria growing in killing conditions over the total recipients was calculated, i.e., the <u>recombination frequency by blind selection</u>.

In all cases, a minimum of eight colonies were verified by PCR with either MFD/SWbeg primers or sacB::attI check R/SWbeg primers.

### Library generation with the cassette gatherer

To create a library from *V. cholerae* N16961 superintegron, we first electroporated this strain A001 with plasmid pB746, which contains *ccdA* and *intI1* (see Fig. 3a). Next, the cells were cultured in the presence of arabinose 0.2% to ensure the production of *ccdA*, before being electroporated with plasmid pB824, which contains the *ccdB::attI1* construction. The next day, at least three transformant colonies were selected and grown in liquid LB medium in the presence of zeocin 50 µg/mL, carbenicillin 100 µg/mL, arabinose 0.2%, and vanillate 25 µM (to induce the expression of the integrase). After 3 hours of incubation, serial 1:10 dilutions of the liquid culture were performed up to a 10^-7^ dilution, and 5µL of each dilution was plated on LB medium supplemented with various antibiotics and reagents. To count the total recipient cells, the culture was plated on zeocin 50 µg/mL, carbenicillin 100 µg/mL, and arabinose 0.2%. To count recombinants, the culture was plated on 50 zeocin µg/mL, carbenicillin 100 µg/mL, glucose 1%, and IPTG 200 µg/mL. The recombination frequency was calculated as the proportion of recombinant CFUs with respect to the total number of recipient CFUs. The cassette content of recombinants was amplified with primers MDF and MRV II. Cassettes were then identified through Sanger sequencing (Macrogen, Korea) using bot primers. The library was stored by extracting plasmids from a pool of ∼10^4^ cells.

### Recombination through natural competence and library generation with the cassette hunter

Although *V. cholerae* N16961 was initially characterized as non-competent due to a frameshift in the *hapR* gene, the phenotype was corrected by introducing *hapR* in a miniTn7^47^. This generated the parental strain of *V. cholerae* N16961 ΔSI, strain B522 (CECT30916), on which all the natural competence experiments in this study are based.

Recombination assays through natural competence were performed with a *tfoX*-inducing competence assay using an adapted protocol from^65^. Strains were inoculated from frozen stocks into liquid LB medium with IPTG 200 µg/mL (in the case of the P_Tac_-controlled *tfoX* plasmid) and grown overnight at 30°C. Overnight cultures were then diluted 1:100 and grown in the appropriate antibiotics, IPTG 200 µg/mL, and arabinose 0.2% for four hours. Cells were washed and resuspended in DASW (Instant Ocean, USA) 0.5x, and 7 µL were inoculated in a final volume of 250 µL DASW (adding 150 µL of chitin slurry in the final assays of the cassette hunter). 2-4 µg of tDNA were added, and cells were incubated in static conditions at 30°C for ∼18 h. As a control of natural competence, 2 µg of tDNA were provided from PCR amplified DNA from strain #135 (Blokesch lab collection) with primers LacZ-FRT1 and LacZ-FRT4. This DNA segment contains the *aph* gene that confers resistance to kanamycin and homology regions targeting the *lacZ* gene. The following day, 1:10 serial dilutions up to 10^-7^ were performed, and 5 µL of each dilution were plated on LB medium supplemented with the appropriate antibiotics and glucose 1% to recover the total recipient bacteria. To recover recombinants or transformants (homologous recombinations) through phenotype selection, cells were plated in either chloramphenicol 2.5 µg/mL + glucose 1% or kanamycin 75 µg/mL + glucose 1%, respectively. To recover recombinants through blind selection, cultures were plated in LB medium (∅NaCl) + sucrose 10% and incubated at 20°C for up to 48h. To characterize the captured cassettes, we performed a colony PCR of the recombinants using the primers sacB::attI check F and sacB::attI check R.

Concerning the transformed DNA samples, the three tested PCR products from the pMBA collection^10,22,39^ were amplified with IntI seq F II and GFP seq R II primers. The tested gDNAs were purified with the DNeasy Blood and Tissue Kit (Qiagen, USA). 4 µg of DNA was supplemented to the competent bacteria in the case of the sedentary chromosomal integron (SCI)-containing samples.

The SCI-generated libraries from the *Vibrio* spp. SCIs were stored by recovering at least 10^3^ CFUs and subsequently sequenced using long-read sequencing. The primers used to amplify the cassette content were sacB::attI check F and sacB::attI check R.

### Library sequencing and analysis

#### Library sequencing and read processing

A PCR of the captured cassettes was performed using primers MFD and MRV II in the case of the cassette gatherer, and sacB::attI check F and R in the case of the cassette hunter. The resulting purified PCR product was sequenced by Plasmidsaurus (USA) using Oxford Nanopore Technology. Reads were processed with a custom pipeline using software packages from Bioconda, CRAN, or Bioconductor public repositories. First, the quality of sequencing reads was assessed using Nanoplot (version 1.42.0). Low-quality reads (filtered to Q >10 and length >100 nt) and adapters were trimmed using NanoFilt (version 2.8.0) and PoreChop (version 0.2.4), respectively. After quality control, the reads were mapped to the reference genomes (*Vibrio cholerae* N16961 (RefSeq GCF_000006745.1), *Vibrio cholerae* SA5Y (RefSeq GCF_003063885.1), *V. cholerae* W6G (RefSeq GCF_013357785.1), *V. vulnificus* NBRC 15645 (RefSeq GCF_002224265.1), *V. mimicus* MB461 (GenBank GCA_ADAF00000000.1), and *V. parahaemolyticus* RIMD 2210633 (RefSeq GCF_000196095.1), using Minimap2 (version 2.28) with the *-ax map-ont* option, allowing for secondary alignments to account for multiple integron recombination events. Mapping coverage was analyzed and visualized using Bedtools (version 2.31.1) and BAMdash (version 0.2.4).

#### Cassette identification and quantification

The structure of the SCIs was annotated with IntegronFinder (version 2.0.6, with options *--local-max --evalue-attc 4 --calin-threshold 1*). In the case of *Vibrio cholerae* N16961, two *attC* sites were manually added (93.02% and 96.72% identity with other *attC* sites from the SI using Geneious software 2025.1.2), as it was previously described that the *V. cholerae* N16961 SI possesses 179 cassettes^66^. Nevertheless, we were only able to identify 178 with this combined approach in our lab strain. Then, cassettes were defined as sequences located between consecutive *attC* sites, or between the *attI* site and the first *attC* site. The SCI annotation was combined with Bakta (software version 1.11, Database v. 5.1) genome annotation to obtain a final updated annotated genome. Read counts per feature (CDS, *attC*, cassette) and features per read were determined using the *featureCounts* function from the Rsubread package (version 2.18.0), with the following options: *isPairedEnd=FALSE, countMultiMappingReads=TRUE, allowMultiOverlap=TRUE*. Data handling, statistical analysis, and plotting were carried out using the R packages circlize, ggplot2, ggplotly, ggpubr, and Biocircos.

### Flow cytometry analysis of fluorescence

Independent colonies were grown overnight in LB liquid medium supplemented with the appropriate antibiotics. The next day, cultures were diluted 1:400 in filtered saline solution (NaCl 0.9%) and fluorescent intensity was measured by flow cytometry using a CytoFLEX-S cytometer (Beckman Coulter, USA). The measurement of each biological replicate is the result of the mean of the fluorescence intensity of 30,000 events per sample. Data processing was performed with Cytexpert software.

### Phage-defense screening

#### Direct screening

To perform the direct phenotypic screening, we used the stored libraries in the cassette hunter chassis, and performed a variant of the “tab” method, as described in ^54^. Briefly, a heavy inoculum from the −80 °C library stock (or empty chassis as a control) was grown overnight in 5 mL LB at 37 °C. Cultures were adjusted to OD_600_ = 1.0, and 0.1 mL was added to a Petri dish. A tenfold dilution series of the phage stock was prepared, and 0.1 mL of each dilution was pipetted onto separate areas of the plate to prevent mixing. Molten LB agar (20 mL, 0.5% agar) was added and briefly mixed to disperse bacteria and phage. Plates were incubated overnight at 37 °C. The next day, bacterial colonies were picked from the plate containing the phage dilution that produced the largest difference in number of colonies between the control and library samples and then streaked onto fresh plates to isolate single colonies. These colonies were used as templates to amplify the cassette content with primers gblock F and sacB pMBA R, which was identified by Sanger sequencing. The cassette was then subcloned into the P_C_W pMBA vector (pD709), linearized using primers IntI R bb and GFP F bb, and the Gibson assembly was transferred into *E. coli* MG1655. After colony PCR verification, plasmids were extracted and electroporated into *V. cholerae* ΔSI Δ*ddm* for further phenotype validation.

#### Indirect screening

To perform the indirect phage screening, we amplified the cassette content of the six generated libraries and cloned them in the P_C_S pMBA vector (pD710), as described above. The Gibson assembly mixes were transferred into *E. coli* MG1655, and at least 10^3^ CFUs were recovered to store the plasmid library. Then, screens were performed as above. Finally, survivors were identified and plasmids re-transformed in a clean pMBA vector and *E. coli* MG1655 host.

#### Phenotype validation

To confirm the phenotype of the potentially newly identified phage-defense systems, phage plaque assays were performed: we first grew overnight cultures of the transformed strains, then adjusted the optical density at 600 nm (OD_600_) to 1.0. Next, we mixed 1 mL of the adjusted cultures with 9 mL of soft LB agar medium (0.4% agar) and poured it onto LB agar plates. After allowing plates to dry for 15 minutes, serial dilutions of each phage were spotted, and plates were incubated overnight at 37 °C. For phage T7, plates were incubated for 8 hours.

#### In silico analysis

We used the AlphaFold3^42^ server from Google DeepMind for oligomerization experiments. Multimers were selected when the interface-predicted template modeling score (ipTM) was > 0.5. Oligomer structures were visualized using ChimeraX. Protein sequences were analyzed with InterProScan^67,68^ via the InterPro resource (European Bioinformatics Institute) to identify conserved domains and predict functional annotations.

### Statistical analysis and reproducibility

Statistical analysis was performed using GraphPad Prism 10.1.1 (GraphPad Software) or R studio. Significant differences were determined using the Mann-Whitney U or Kruskal-Wallis tests, with multiple comparisons corrected by Dunnett’s test. A significance level of 0.05 was applied in all cases. Significance levels were indicated as follows: *p* ≥ 0.05 (ns), *p* < 0.05 (*), *p* < 0.01 (**), *p* < 0.001 (***), and *p* < 0.0001 (****). Correlations were calculated with Spearman algorithm in ggscatter package or Prism 10.1.1.

All experimental data are representative of at least three independent biological repeats.

## Data availability

All raw sequencing and analysis data generated for this project can be found on Zenodo Data Repository: https://doi.org/10.5281/zenodo.18864999. Library analysis was performed using custom R scripts, described in the GitHub Repository: https://github.com/mredrejo/cassette_gatherer_hunter.

## ACKOWLEGDMENTS

We thank Dr. José R. Penadés for the constructive comments on the manuscript, Dr. Ankur Dalia for providing materials, and Dr. David Adams for helpful discussion.

This work was supported by the European Research Council (ERC) through a Starting Grant [ERC grant no. 803375-KRYPTONINT]; Ministerio de Ciencia, Innovación y Universidades [BIO2017-85056-P, PID2020-117499RB-100]; J.A.E. was supported by the Atracción de Talento Program of the Comunidad de Madrid [2016-T1/BIO-1105 and 2020-5A/BIO-19726]; F.T.R. was supported by the Portuguese Fundação para Ciência e a Tecnologia [SFRH/BD/144108/2019] and with an EMBO Scientific Exchange Grant [10279]; A.M. is supported by the Universidad Complutense de Madrid; P.B. was supported by the Juan de la Cierva program [FJC 2020-043017-I]; R.L.-I. was supported by Ministerio de Ciencia e Innovación and the EU “NextGenerationEU”/PRTR” [RYC2021-034768-I, CNS2023-145397]; M.R.R. was supported by Ministerio de Ciencia e Innovación and the European Regional Development Fund [MCIN/AEI/10.13039/501100011033, PID2021-123403NB-I00]; M.B. was supported by intramural funding by EPFL.

## AUTHOR CONTRIBUTIONS

F.T.R.: Conceptualization, methodology, formal analysis, validation, writing—original draft, review and editing; A.C.: Conceptualization, methodology, analysis and validation; A.P.: Methodology, analysis and validation; P.B.: Methodology, analysis and validation; E.V.: Methodology, analysis and validation; R.L.-I.: Methodology, analysis and validation; M.R.R.: Methodology, analysis and validation; M.B.: Conceptualization, writing—review and editing; J.A.E.: Conceptualization, methodology, formal analysis, validation, writing—original draft, review and editing.

## COMPETING INTERESTS

The *V. cholerae* ΔSI strain is covered by patent ES2970040B2 (international extension WO2025104363). The *ccdB*/*ccdA* plasmid-based tool is covered by patent ES2969666B2. The *sacB* chromosome-based tool is covered by patent ES2991744B2 (international extension WO2025238278).

---

**Extended Data Figure 1.**
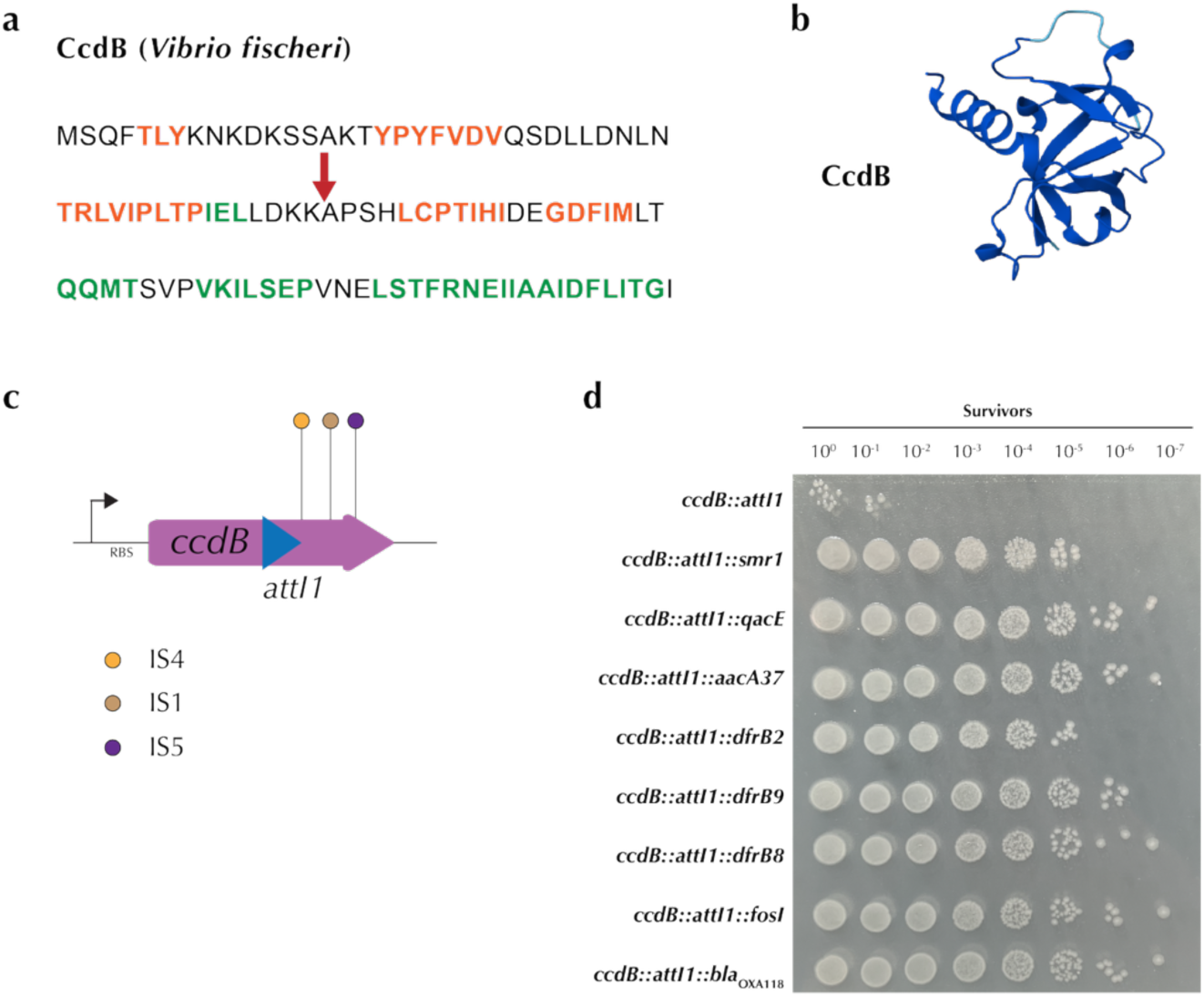
**a)** Chosen location to introduce the *attI1* site in *ccdB* from *Vibrio fischeri*. Orange represents β-sheets and green represents α-helices. These features were predicted using Phyre2. The red arrow illustrates the locations where the *attI1* site was introduced. **b)** Predicted structure of wild-type CcdB protein from *V. fischeri*. Models were generated using AlphaFold3^42^; the blue-to-red scale indicates high to low confidence in folding prediction. pTM = 0.9. **c)** Schematic representation of the escape mutants encountered in the killing assays of *ccdB::attI1*. Colonies were amplified by PCR and mutations identified by Sanger-sequencing (n=30). **d)** Killing assays of *ccdB::attI1::cassette*. The *E. coli* strains contain the various plasmidic constructs mimicking the outcome of a recombination event. These strains were plated in 5 µL spots in dilutions up to 10^-7^, in killing conditions.

**Extended Data Figure 2.**
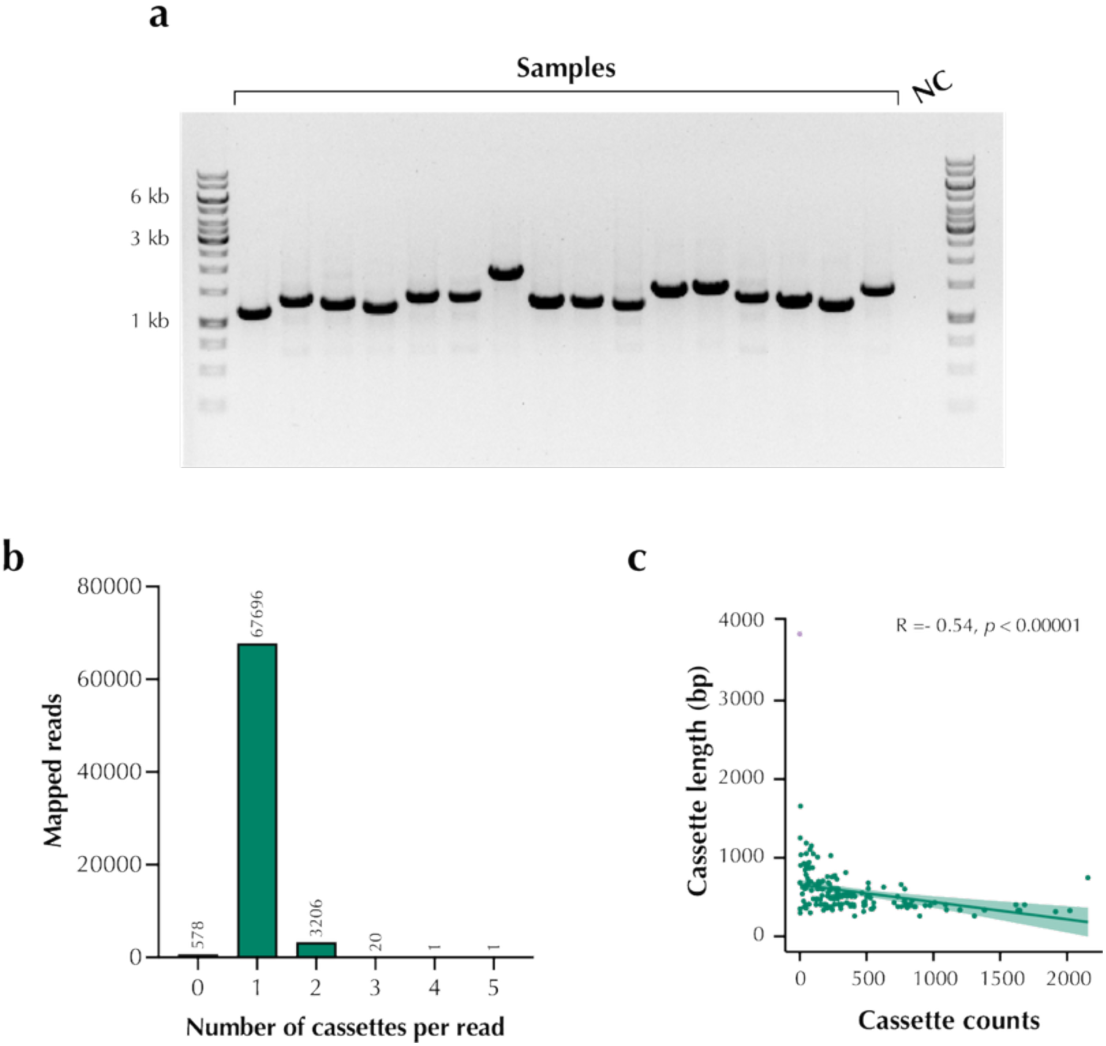
**a)** Example of a colony PCR result of the captured cassettes in the plasmid vector. The expected size of the empty *ccdB::attI1* would be 667 bp. NC, PCR negative control. **b)** Number of reads that mapped to 0-5 cassettes. **c)** Correlation of the cassette length and the number of counts of the same cassette. Spearman R and *p*-value are shown.

**Extended Data Figure 3.**
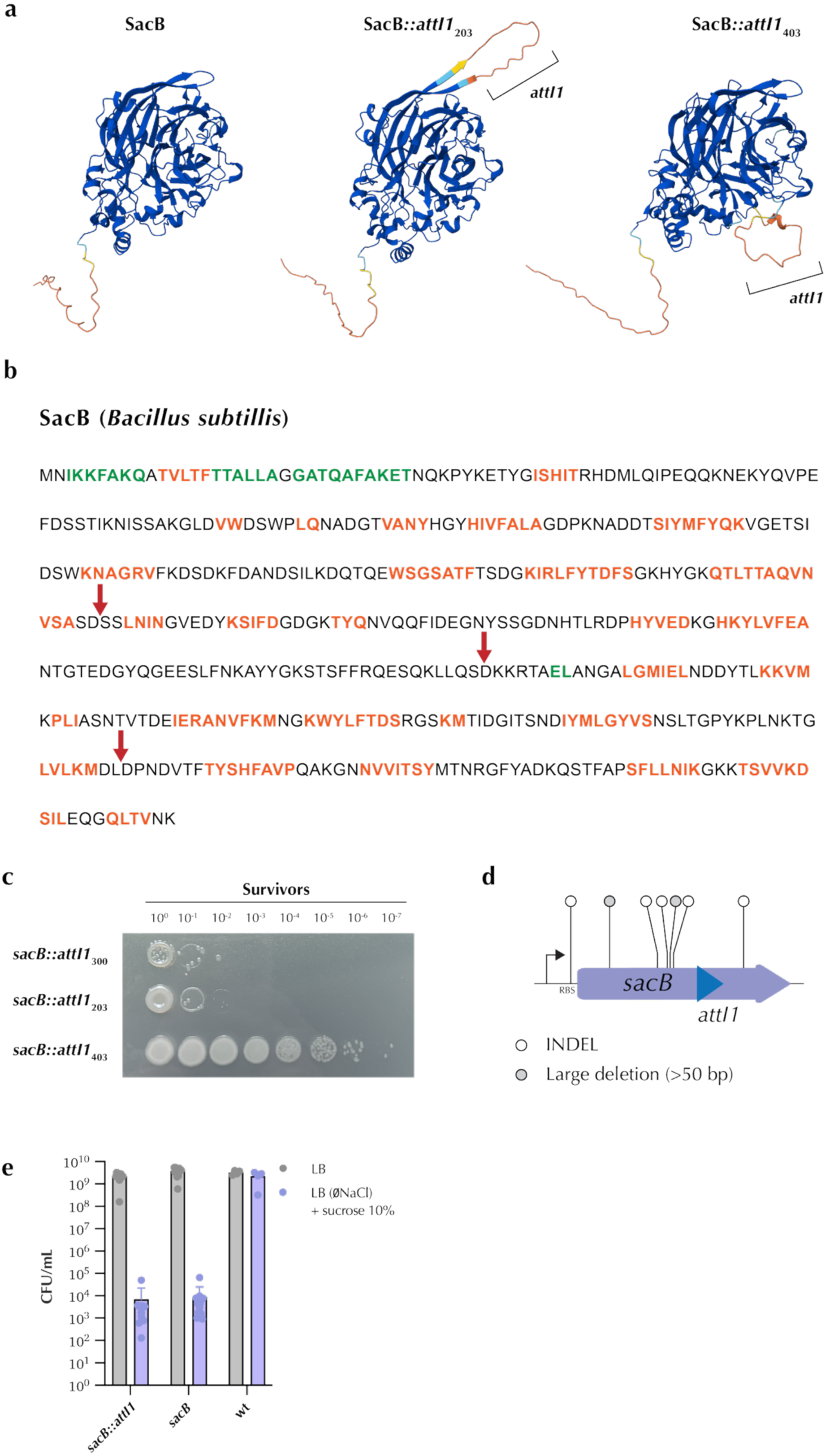
**a)** Predicted structures of wild-type SacB protein from *Bacillus subtilis* and SacB::*attI1*_203_ and SacB::*attI1*_403_, highlighting the integration *attI1* site and altered folding. Models were generated using AlphaFold3^42^, blue-to-red scale indicates high to low confidence in folding prediction. pTM = 0.92, 0.9, and 0.89, respectively. **b)** Chosen locations to introduce the *attI1* site in *sacB* from *B. subtilis*. Orange represents β-sheets and green represents α-helices. These features were predicted using Phyre2. Red arrows illustrate the locations where the *attI1* site was introduced. **c)** Survivors in a killing assay testing the three SacB::*attI1* versions. The three constructs were plated in 5µL spots in dilutions up to 10^-7^, in killing conditions in LB (∅NaCl) + sucrose 10%. **d)** Schematic representation of the escape mutants encountered in the killing assays of *sacB::attI1*. Colonies were amplified by PCR and mutations identified by Sanger-sequencing (n=10). **e)** Cell counts of the killing assays of the *sacB*-based platform cloned in *V. cholerae* N16961 ΔSI compared to the WT gene. The WT strain (*sacB* free) was used as a control for growth in the different tested media. Cells were grown in either survival conditions (standard LB medium) or killing conditions (LB (∅NaCl) + sucrose 10%).

**Extended Data Figure 4.**
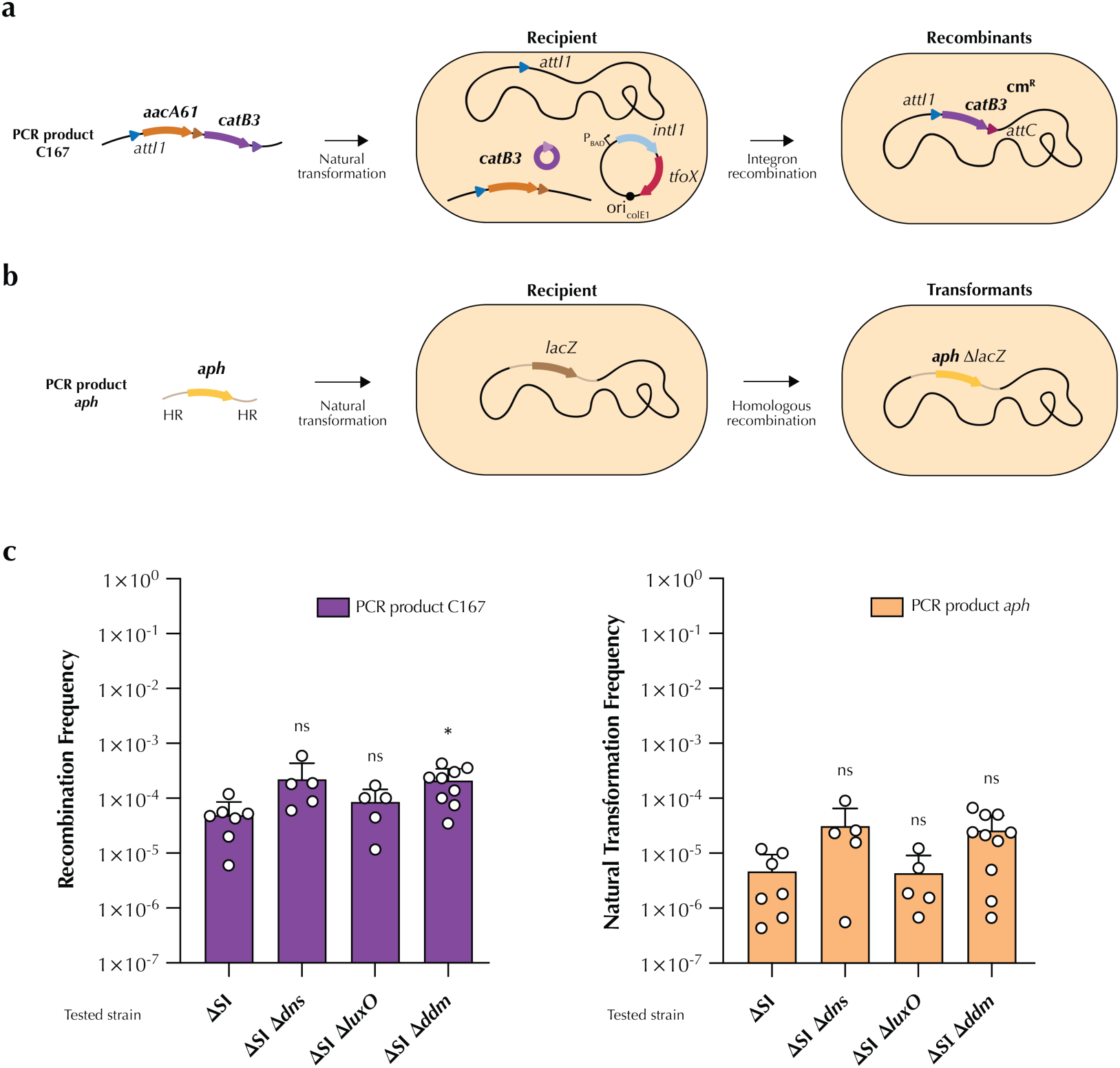
**a)** Schematic representation of the recombination assay. Recipient cells were transformed with a PCR-amplified integron array upon arabinose-mediated *tfoX* and *intI1* induction. The different *V. cholerae* ΔSI Δ*ddm* mutants contain an *attI1* site in the chromosome. The capture of the second cassette (*catB3*) was selected by phenotype, i.e., chloramphenicol resistance. **b)** Schematic representation of the classical natural transformation assay. Above, a *V. cholerae* recipient is depicted with the *lacZ* gene and its adjacent regions. The transforming DNA corresponds to a kanamycin resistance gene (*aph*) and its promoter, flanked by adjacent regions of the target (in light brown), called homology regions (HR). After transformation, the *aph* gene substitutes the *lacZ* gene by homologous recombination, allowing for the selection of transformants. **c)** Recombination and transformation frequencies are given on the Y-axis and were calculated as the ratio of the number of CFUs resistant to chloramphenicol or kanamycin versus the total number of CFUs. Bar charts show the mean of the recombination/transformation frequency ± s.d. from several biological replicates (individual dots). Statistical significance between each mutant and the ΔSI control was determined using the Kruskal-Wallis test with Dunnett’s correction for multiple comparisons; ns, not significant, *, *p* < 0.05.

**Extended Data Figure 5.**
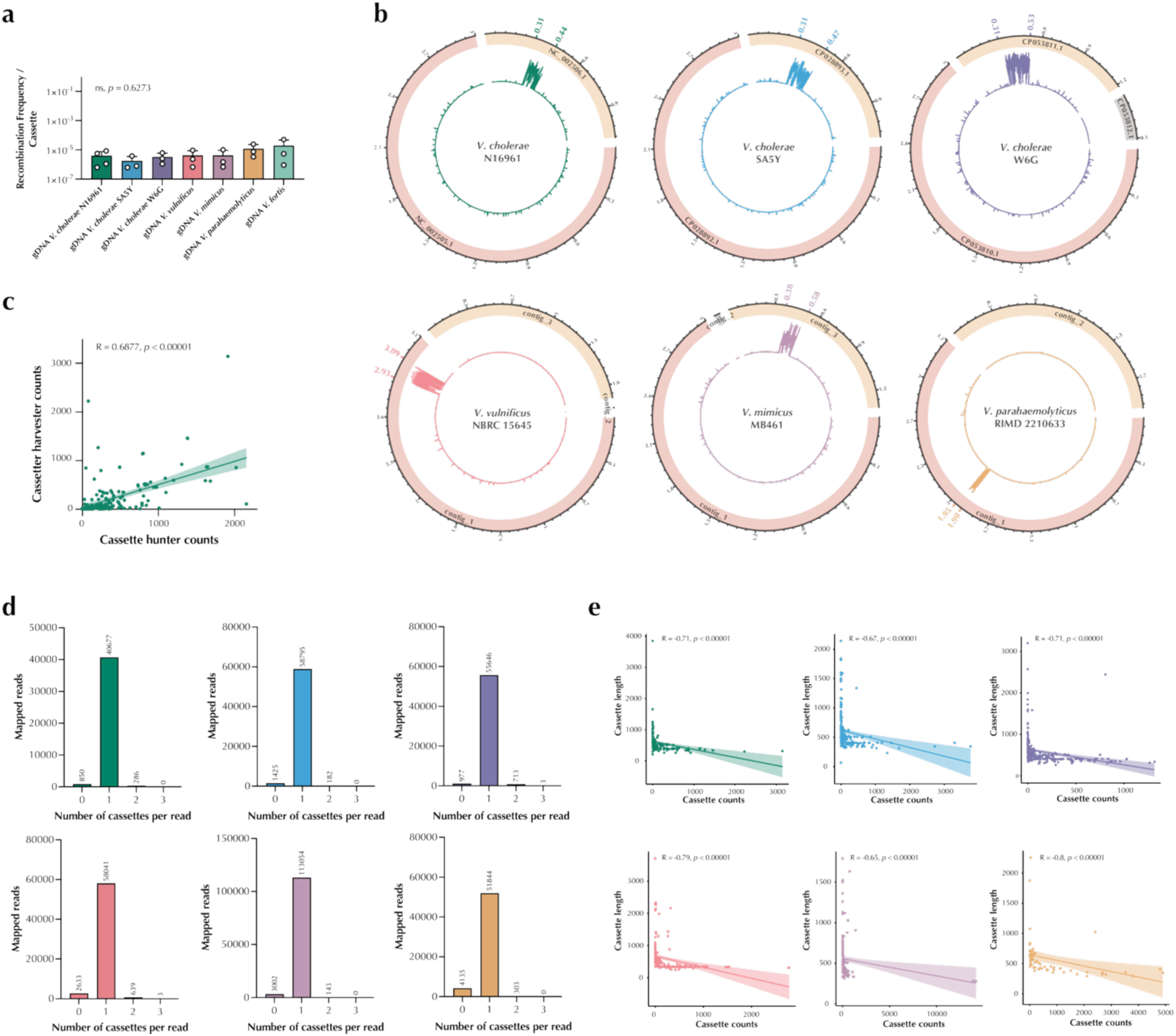
**a)** Recombination frequency per cassette of each genomic sample tested. Statistical significance between samples was determined using the Kruskal-Wallis test with Dunnett’s correction for multiple comparisons; ns, not significant. **b)** Average coverage per kilobase (colored plots) given in log10 reads of the contigs of each *Vibrio* spp. genome. SCI positions are highlighted in bold. **c)** Correlation between the cassette counts retrieved on the cassette gatherer tool and the cassette hunter tool for the *V. cholerae* N16961 SCI. **d)** Number of reads that mapped to 0-5 cassettes. **e)** Correlation of the cassette length and the number of counts of the same cassette for each library generated. Spearman R and p-value are shown.

**Extended Data Figure 6.**
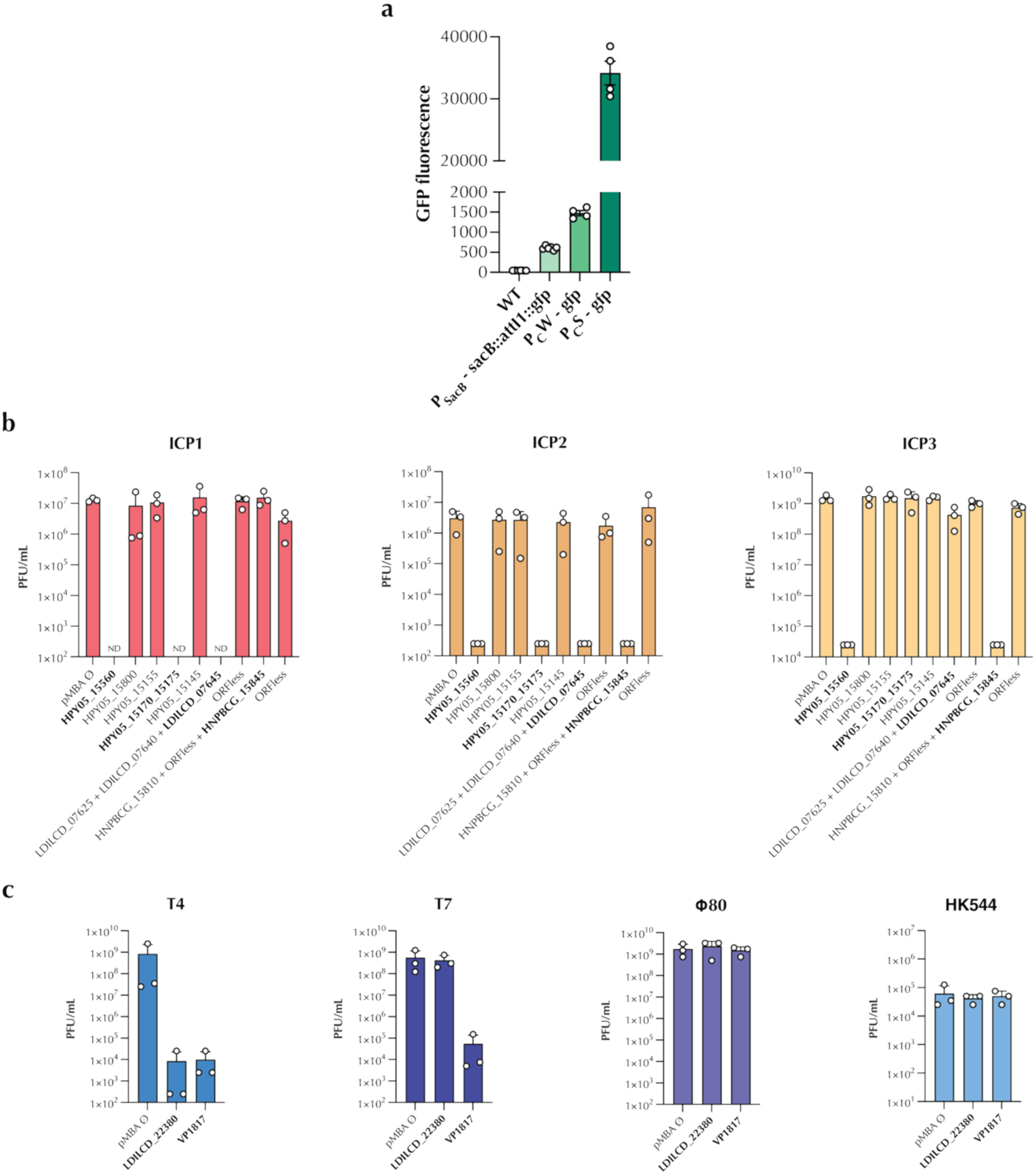
**a)** GFP fluorescence intensity (FITC-A, arbitrary units) measured by flow cytometry of three different promoters controlling the *gfp* gene in *V. cholerae* N16961. WT represents the empty strain, as a no-fluorescent control. Dots represent biological replicates. **b)** Plaque-forming units (PFU) per mL of tested integron cassettes against Vibriophages ICP1, ICP2, and ICP3. **c)** Plate-forming units (PFU) per mL of tested integron cassettes against *E. coli* phages T4, T7, Φ80, and HK544.

## Notes

### Competing Interest Statement

The V. cholerae ∆SI strain is covered by patent ES2970040B2 (international extension WO2025104363) published on 16/01/2025. The ccdB/ccdA plasmid-based tool is covered by patent ES2969666B2 published on 16/01/2025. The sacB chromosome-based tool is covered by patent ES2991744B2 (international extension WO2025238278) published on 23/04/2025.

## REFERENCES

1. Escudero, J., Loot, C., Nivina, A. & Mazel, D. The Integron: Adaptation On Demand. Microbiol. Spectr. 13, MDNA3-0019–2014 (2015).

2. Mazel, D. Integrons: Agents of bacterial evolution. Nat. Rev. Microbiol. 4, 608–620 (2006).

3. Mitsuhashi, S., Harada, K., Hashimoto, H. & Egawa, R. On the drug-resistance of enteric bacteria. 4. Drug-resistance of Shigella prevalent in Japan. Jpn. J. Exp. Med. 31, 47–52 (1961).

4. Kieffer, N. et al. Mobile integrons encode phage defense systems. Science 388, eads0915 (2025).

5. Darracq, B. et al. Sedentary chromosomal integrons as biobanks of bacterial antiphage defense systems. Science 388, eads0768 (2025).

6. Getz, L. J., Fairburn, S. R., Vivian Liu, Y., Qian, A. L. & Maxwell, K. L. Integrons are anti-phage defence libraries in Vibrio parahaemolyticus. Nat. Microbiol. 10, 724–733 (2025).

7. Boucher, Y., Labbate, M., Koenig, J. E. & Stokes, H. W. Integrons: mobilizable platforms that promote genetic diversity in bacteria. Trends Microbiol. 15, 301–309 (2007).

8. Cury, J., Jové, T., Touchon, M., Néron, B. & Rocha, E. P. Identification and analysis of integrons and cassette arrays in bacterial genomes. Nucleic Acids Res. 44, 4539–4550 (2016).

9. Partridge, S. R., Tsafnat, G., Coiera, E. & Iredell, J. R. Gene cassettes and cassette arrays in mobile resistance integrons. FEMS Microbiol. Rev. 33, 757–784 (2009).

10. Hipólito, A., García-Pastor, L., Vergara, E., Jové, T. & Escudero, J. A. Profile and resistance levels of 136 integron resistance genes. Npj Antimicrob. Resist. 1, 1–12 (2023).

11. Gillings, M. R. Class 1 integrons as invasive species. Curr. Opin. Microbiol. 38, 10–15 (2017).

12. Zhu, Y.-G. et al. Microbial mass movements. Science 357, 1099–1100 (2017).

13. Cambray, G., Guerout, A.-M. & Mazel, D. Integrons. Annu. Rev. Genet. 44, 141–166 (2010).

14. Blanco, P. et al. Chromosomal integrons are genetically and functionally isolated units of genomes. Nucleic Acids Res. 1–17 (2024) 10.1093/nar/gkae866.

15. Buongermino Pereira, M., et al. A comprehensive survey of integron-associated genes present in metagenomes. BMC Genomics 21, 495 (2020).

16. Mazel, D., Dychinco, B., Webb, V. A. & Davies, J. A distinctive class of integron in the Vibrio cholerae genome. Science 280, 605–608 (1998).

17. Escudero, J. A. & Mazel, D. Genomic Plasticity of Vibrio cholerae. Int. Microbiol. 20, 138–148 (2017).

18. Rowe-Magnus, D. A., Guerout, A. M. & Mazel, D. Bacterial resistance evolution by recruitment of super-integron gene cassettes. Mol. Microbiol. 43, 1657–1669 (2002).

19. Nunes-Düby, S. E., Kwon, H. J., Tirumalai, R. S., Ellenberger, T. & Landy, A. Similarities and differences among 105 members of the Int family of site-specific recombinases. Nucleic Acids Res. 26, 391–406 (1998).

20. Recchia, G. D., Stokes, H. W. & Hall, R. M. Characterisation of specific and secondary recombination sites recognised by the integron DNA integrase. Nucleic Acids Res. 22, 2071–2078 (1994).

21. Gravel, A., Fournier, B. & Roy, P. H. DNA complexes obtained with the integron integrase Intl1 at the attl1 site. Nucleic Acids Res. 26, 4347–4355 (1998).

22. Carvalho, A. et al. The expression of integron arrays is shaped by the translation rate of cassettes. Nat. Commun. 15, 9232 (2024).

23. Jové, T., Da Re, S., Denis, F., Mazel, D. & Ploy, M. C. Inverse correlation between promoter strength and excision activity in class 1 integrons. PLoS Genet. 6, e1000793 (2010).

24. Lévesque, C., Brassard, S., Lapointe, J. & Roy, P. H. Diversity and relative strength of tandem promoters for the antibiotic-resistance genes of several integrons. Gene 142, 49–54 (1994).

25. Recchia, G. D. & Hall, R. M. Gene cassettes: A new class of mobile element. Microbiology 141, 3015–3027 (1995).

26. Stokes, H. W., O’Gorman, D. B., Recchia, G. D., Parsekhian, M. & Hall, R. M. Structure and function of 59-base element recombination sites associated with mobile gene cassettes. Mol. Microbiol. 26, 731–745 (1997).

27. Bouvier, M., Ducos-Galand, M., Loot, C., Bikard, D. & Mazel, D. Structural features of single-stranded integron cassette attC sites and their role in strand selection. PLoS Genet. 5, e1000632 (2009).

28. Loot, C., Bikard, D., Rachlin, A. & Mazel, D. Cellular pathways controlling integron cassette site folding. EMBO J. 29, 2623–2634 (2010).

29. MacDonald, D., Demarre, G., Bouvier, M., Mazel, D. & Gopaul, D. N. Structural basis for broad DNA-specificity in integron recombination. Nature 440, 1157–1162 (2006).

30. Nivina, A., Escudero, J. A., Vit, C., Mazel, D. & Loot, C. Efficiency of integron cassette insertion in correct orientation is ensured by the interplay of the three unpaired features of attC recombination sites. Nucleic Acids Res. 44, 7792–7803 (2016).

31. Escudero, J. A. et al. Unmasking the ancestral activity of integron integrases reveals a smooth evolutionary transition during functional innovation. Nat. Commun. 7, 10937 (2016).

32. Rowe-Magnus, D. A., Guerout, A. M., Biskri, L., Bouige, P. & Mazel, D. Comparative analysis of superintegrons: engineering extensive genetic diversity in the Vibrionaceae. Genome Res. 13, 428–442 (2003).

33. Sandoval-Quintana, E., Lauga, B. & Cagnon, C. Environmental integrons: the dark side of the integron world. Trends Microbiol. 31, 432–434 (2023).

34. Lévesque, C., Piche, L., Larose, C. & Roy, P. H. PCR mapping of integrons reveals several novel combinations of resistance genes. Antimicrob. Agents Chemother. 39, 185–191 (1995).

35. White, P. A., Iver, C. J. M. C. & Rawlinson, W. D. Integrons and Gene Cassettes in the Enterobacteriaceae Downloaded from http://aac.asm.org/ on December 30xlink:href=" 2015 by TERKKO NATIONAL LIBRARY OF HEALTH SCIENCES. Antimicrob. Agents Chemother. 45, 2658–2661 (2001).

36. White, P. A., McIver, C. J., Deng, Y. M. & Rawlinson, W. D. Characterisation of two new gene cassettes, aadA5 and dfrA17. FEMS Microbiol. Lett. 182, 265–269 (2000).

37. Stokes, H. W. et al. Gene cassette PCR: sequence-independent recovery of entire genes from environmental DNA. Appl. Environ. Microbiol. 67, 5240–5246 (2001).

38. Ghaly, T. M., Tetu, S. G. & Gillings, M. R. Predicting the taxonomic and environmental sources of integron gene cassettes using structural and sequence homology of attC sites. *Commun*. Biol. 4, 1–8 (2021).

39. Hipólito, A. et al. The expression of aminoglycoside resistance genes in integron cassettes is not controlled by riboswitches. Nucleic Acids Res. 50, 8566–8579 (2022).

40. Biskri, L., Bouvier, M., Guérout, A.-M., Boisnard, S. & Mazel, D. Comparative Study of Class 1 Integron and Vibrio cholerae Superintegron Integrase Activities. J. Bacteriol. 187, 1740–1750 (2005).

41. López-Igual, R., Bernal-Bayard, J., Rodríguez-Patón, A., Ghigo, J.-M. & Mazel, D. Engineered toxin–intein antimicrobials can selectively target and kill antibiotic-resistant bacteria in mixed populations. Nat. Biotechnol. 37, 755–760 (2019).

42. Abramson, J. et al. Accurate structure prediction of biomolecular interactions with AlphaFold 3. Nature 630, 493–500 (2024).

43. Meyer, A. J., Segall-Shapiro, T. H., Glassey, E., Zhang, J. & Voigt, C. A. Escherichia coli ‘Marionette’ strains with 12 highly optimized small-molecule sensors. Nat. Chem. Biol. 15, 196–204 (2019).

44. Bouvier, M., Demarre, G. & Mazel, D. Integron cassette insertion: a recombination process involving a folded single strand substrate. EMBO J. 24, 4356–4367 (2005).

45. Rodríguez-Beltrán, J. et al. Genetic dominance governs the evolution and spread of mobile genetic elements in bacteria. Proc. Natl. Acad. Sci. 117, 15755–15762 (2020).

46. Loot, C. et al. Differences in Integron Cassette Excision Dynamics Shape a Trade-Off between Evolvability and Genetic Capacitance. mBio 8, e02296–16 (2017).

47. Meibom, K. L., Blokesch, M., Dolganov, N. A., Wu, C.-Y. & Schoolnik, G. K. Chitin Induces Natural Competence in *Vibrio cholerae*. Science 310, 1824–1827 (2005).

48. Gay, P., Le Coq, D., Steinmetz, M., Berkelman, T. & Kado, C. I. Positive selection procedure for entrapment of insertion sequence elements in gram-negative bacteria. J. Bacteriol. 164, 918–921 (1985).

49. Blomfield, I. C., Vaughn, V., Rest, R. F. & Eisenstein, B. I. Allelic exchange in Escherichia coli using the Bacillus subtilis sacB gene and a temperature-sensitive pSC101 replicon. Mol. Microbiol. 5, 1447–1457 (1991).

50. Vit, C. et al. Cassette recruitment in the chromosomal Integron of *Vibrio cholerae*. Nucleic Acids Res. 49, 5654–5670 (2021).

51. Suckow, G., Seitz, P. & Blokesch, M. Quorum Sensing Contributes to Natural Transformation of Vibrio cholerae in a Species-Specific Manner. J. Bacteriol. 193, 4914–4924 (2011).

52. Blokesch, M. & Schoolnik, G. K. The extracellular nuclease Dns and its role in natural transformation of Vibrio cholerae. J. Bacteriol. 190, 7232–7240 (2008).

53. Jaskólska, M., Adams, D. W. & Blokesch, M. Two defence systems eliminate plasmids from seventh pandemic Vibrio cholerae. Nature 604, 323–329 (2022).

54. Vassallo, C. N., Doering, C. R., Littlehale, M. L., Teodoro, G. I. C. & Laub, M. T. A functional selection reveals previously undetected anti-phage defence systems in the E. coli pangenome. Nat. Microbiol. 7, 1568–1579 (2022).

55. Takahashi, H., Coppo, A., Manzi, A., Martire, G. & Pulitzer, J. F. Design of a system of conditional lethal mutations (tab/k/com) affecting protein-protein interactions in bacteriophage T4-infected Escherichia coli. J. Mol. Biol. 96, 563–578 (1975).

56. Maffei, E. et al. Systematic exploration of Escherichia coli phage–host interactions with the BASEL phage collection. PLOS Biol. 19, e3001424 (2021).

57. Tesson, F. et al. Systematic and quantitative view of the antiviral arsenal of prokaryotes. Nat. Commun. 13, 2561 (2022).

58. Payne, L. J. et al. PADLOC: a web server for the identification of antiviral defence systems in microbial genomes. Nucleic Acids Res. 50, W541–W550 (2022).

59. Rowe-Magnus, D. A. Integrase-directed recovery of functional genes from genomic libraries. Nucleic Acids Res. 37, 31–33 (2009).

60. Bikard, D., Julié-Galau, S., Cambray, G. & Mazel, D. The synthetic integron: An in vivo genetic shuffling device. Nucleic Acids Res. 38, e153 (2010).

61. Nivina, A. et al. Structure-specific DNA recombination sites: Design, validation, and machine learning–based refinement. Sci. Adv. 6, eaay2922 (2020).

62. Gibson, D. G. et al. Enzymatic assembly of DNA molecules up to several hundred kilobases. Nat. Methods 6, 343–345 (2009).

63. Stutzmann, S. & Blokesch, M. Comparison of chitin-induced natural transformation in pandemic *Vibrio cholerae* O1 El Tor strains. Environ. Microbiol. 22, 4149–4166 (2020).

64. Meibom, K. L. et al. The Vibrio cholerae chitin utilization program. Proc. Natl. Acad. Sci. U. S. A. 101, 2524–2529 (2004).

65. Chlebek, J. L. et al. PilT and PilU are homohexameric ATPases that coordinate to retract type IVa pili. PLOS Genet. 15, e1008448 (2019).

66. Rowe-Magnus, D. A., Guérout, A. M. & Mazel, D. Super-integrons. Res. Microbiol. 150, 641–651 (1999).

67. Paysan-Lafosse, T. et al. The Pfam protein families database: embracing AI/ML. Nucleic Acids Res. 53, D523–D534 (2025).

68. Blum, M. et al. InterPro: the protein sequence classification resource in 2025. Nucleic Acids Res. 53, D444–D456 (2025).

